# Recruitment orders underlying binocular coordination of eye position and velocity in the larval zebrafish hindbrain

**DOI:** 10.1101/754226

**Authors:** Christian Brysch, Claire Leyden, Aristides B. Arrenberg

**Author notes:** Correspondence should be addressed to A.B.A.

## Abstract

**Background:** The oculomotor integrator (OI) in the vertebrate hindbrain transforms eye velocity input into persistent position coding output, which plays a crucial role in retinal image stability. For a mechanistic understanding of the integrator function and eye position control, knowledge about the tuning of the OI and other oculomotor nuclei is needed. Zebrafish are increasingly used to study integrator function and sensorimotor circuits, yet the precise neuronal tuning to motor variables remains uncharacterized.

**Results:** Here, we recorded cellular calcium signals while evoking monocular and binocular optokinetic eye movements at different slow-phase eye velocities. Our analysis reveals the anatomical distributions of motoneurons and internuclear neurons in the nucleus abducens as well as those of oculomotor neurons in caudally adjacent hindbrain volumes. Each neuron is tuned to eye position and/or velocity to variable extents and is only activated after surpassing particular eye position and velocity thresholds. While the abducens (rhombomeres 5/6) mainly codes for eye position, in rhombomeres 7/8 a velocity-to-position coding gradient exists along the rostro-caudal axis, which likely corresponds to the velocity and position storage mechanisms. Position encoding neurons are recruited at eye position thresholds distributed across the behavioral dynamic range, while velocity encoding neurons have more centered firing thresholds for velocity. In the abducens, neurons coding exclusively for one eye intermingle with neurons coding for both eyes. Many of these binocular neurons are preferentially active during conjugate eye movements, which represents a functional diversification in the final common motor pathway.

**Conclusions:** We localized and functionally characterized the repertoire of oculomotor neurons in the zebrafish hindbrain. Our findings provide evidence for a mixed but task-specific binocular code and suggest that generation of persistent activity is organized along the rostro-caudal axis in the hindbrain.

## Background

The oculomotor system is responsible for moving the eyes in vertebrates and is highly conserved across species. Zebrafish are increasingly used to improve our understanding of the oculomotor population code and eye movement control [1]–[6].

The oculomotor system for horizontal eye movements consists of multiple elements (Fig. 1a). It is responsible for generating and maintaining stable eye positions as well as eye movements during saccades, optokinetic and vestibulo-ocular reflexes (OKR, VOR), and other behaviours. The lateral and medial rectus (LR, MR), which represent the extraocular eye muscles responsible for horizontal eye movements, are controlled by motoneurons (MN) in the nucleus abducens (ABN) and the oculomotor nucleus (OMN), respectively. The OMN MNs are activated by internuclear neurons (INN) residing in the contralateral ABN. The ABN receives input from a range of structures such as the burst (B) system for driving saccades, the horizontal eye velocity-to-position neural integrator (termed oculomotor integrator, OI) for maintaining eye positions (P), and the velocity storage mechanism (VSM) associated with slow eye velocities (V) during optokinetic and vestibular responses.

**Figure 1:**
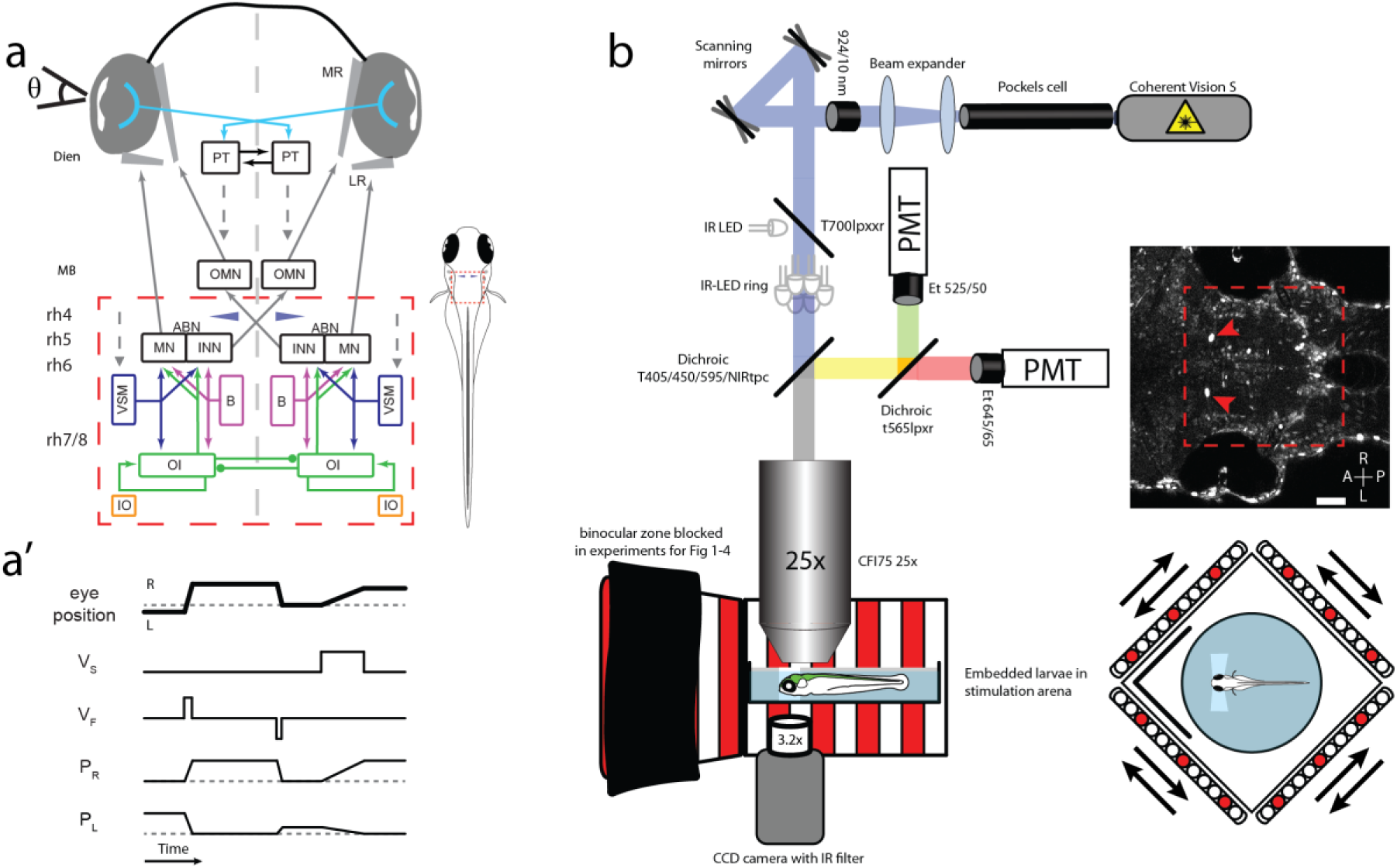
Setup & Circuit overview. **a:** Circuit schematic for horizontal eye movements. Red dashed rectangle represents imaged brain area, blue cones show location of Mauthner cells. ABN: abducens nucleus; B: burst neurons; Dien: diencephalon; INN: internuclear neurons; IO: inferior olive; LR: lateral rectus; MB: midbrain; MN: moto neurons; MR: medial rectus; OMN: nucleus oculomotorius; OI: oculomotor integrator; PT: pretectum; rh 4-8: rhombomeres 4-8; VSM: velocity storage mechanism; Θ: eye position. Dashed arrows indicate direct or indirect inputs from upstream visual brain areas. **a’:** Simplified schematic response profiles for hindbrain oculomotor neurons during eye position changes. Dashed line represents an eye position or velocity of 0. L: left PL/R: Position coding neurons left/right, note that P_L_ and P_R_ have different firing thresholds; R: right; V_F_: fast (burst) velocity neurons; V_S_: slow velocity neurons. **b:** Schematic of microscopy setup. Agarose-embedded zebrafish larvae were visually stimulated, while eye movements were recorded from below and cellular calcium signals were recorded from above via a two-photon microscope. Setup not drawn to scale, binocular zone excluded for experiment with monocular stimulation only, scale bar 50 µm, red dashed rectangle represents imaged brain area, red arrows show GCaMP expression in the nuclei of the Mauthner cells, which served as a landmark (blue cones in a and in cell maps). A: anterior; L: left; P: posterior; PMT: photomultiplier tubes R: right.

The oculomotor integrator is of particular interest, as its persistent firing and dynamic integration of inputs manifest a short-term memory of eye position. It mathematically integrates eye velocity inputs in order to generate a neural representation of eye position via persistent firing [7], [8]. Its mechanisms of operation [9]–[11] are not fully understood and could provide insights into memory functions of other, higher, brain areas as well. The OI neurons in zebrafish are functionally heterogeneous and their differential function is likely related to the mechanism of integration. The zebrafish OI is located in hindbrain rhombomeres 7 and 8 and is organized internally along both the rostro-caudal and dorsal-ventral axes, resulting in a gradient of neuronal persistence times [12]. Neurotransmitter identities as well as axonal projection patterns have been characterized previously ([13]– [15]). In theoretical models of integration mechanisms [9]–[11], [16], [17], the existing recruitment order of integrator neurons is crucial: each neuron carries an eye position threshold and once surpassed, the firing rate is linearly related to the eye position in the ON direction [18]–[20].

In the cat and primate brain, the OI is located in two nuclei, the nucleus prepositus hypoglossi (NPH) and the medial vestibular nucleus (MVN). It contains position coding neurons, which in addition encode saccadic eye velocity to variable extents ([19], [20]). In the goldfish OI (termed Area I in goldfish) position neurons typically also encode saccadic velocity [18].

The velocity storage mechanism is a second short-term memory system in the oculomotor hindbrain, which is charged by vestibular or optic flow stimulation via vestibular nuclei and the pretectum/accessory optic system. It supports retinal and global image stabilization and maintains the eye velocity for a certain time after cessation of stimulus movement in an after-response. While the monkey NPH has been reported to encode eye/head velocity during vestibular stimulation [19] as well, in goldfish such head velocity signals are restricted to an anatomical region termed Area II, which is located rostral to the OI [21]–[23]. The low-velocity encoding neurons have not been functionally identified in zebrafish yet [but see anatomical regions in [2], [22]]. Zebrafish readily generate slow-phase optokinetic reposes and therefore velocity encoding neurons are needed. However, the VSM is still immature in developing larvae: velocity is only stored for very brief periods of time – if at all [24], [25].

In summary, the differential eye position and velocity tuning of zebrafish hindbrain neurons is still elusive but crucial for understanding the functional architecture of the OI and other oculomotor nuclei. Here, we employ stimulus protocols designed to measure eye position and eye velocity encoding independently and reveal an anatomical velocity-to-position gradient in rhombomeres 7 and 8 as well as recruitment orders for eye position and eye velocity during the slow phase of the OKR.

In addition to the position/velocity tuning, we characterize the ocular tuning in this study. Since vertebrates possess two eyes, the drive for each eye needs to be binocularly coordinated to facilitate stable perception of the whole visual field. This binocular coordination is a readily observable feature in human and zebrafish oculomotor behaviour: most of the time both eyes move in the same direction with the same amplitude. Historically, two different mechanism have been suggested: The two eyes could receive conjugate commands to move together “as two horses on the one rein” (Hering’s hypothesis), or each eye could be controlled independently so that binocular coordination would need to be learned (Helmholtz’ hypothesis, [26], [27]). It remains uncertain how binocular coordination is implemented, with the likelihood that a full explanation contains elements of both theories [28], [29]. Here, we employ monocular and binocular stimulation protocols to drive conjugate and monocular eye movements while measuring neuronal activity. We present evidence for a mixed mono-/binocular code in the hindbrain. Within the abducens nucleus different neurons are recruited preferentially during binocular versus monocular optokinetic responses, which represents a deviation from a strict final common motor pathway.

## Results

### Zebrafish hindbrain neurons group into distinct mono- and binocular clusters

To localize and functionally characterize hindbrain neurons active during oculomotor behaviour, we stimulated larvae with patterns of moving gratings to elicit optokinetic responses while measuring GCaMP6f calcium signals in individual neurons (Fig. 1a-b).

Zebrafish show a high degree of binocular coordination: most of the time, the eyes are moved in a conjugate fashion with the notable exception of convergence during prey capture and spontaneous monocular saccades [[30], own observations]. In order to assess the binocular coordination within the oculomotor system and to identify the location of internuclear neurons (INNs) and other structures, we applied a stimulus protocol geared to decouple both eyes and reduce the gain of the non-stimulated eye to <0.1 by showing a moving grating to the stimulated and a stable grating to the non-stimulated eye [24]. This enabled us to classify neurons according to their innervated eye(s) based on their response profile. The stimulus consisted of stimuli driving primarily monocular and conjugate eye movements, respectively. The large decorrelation of left and right eye movements enabled us to classify the monocular or binocular coding of each neuron (Fig. 2a-b’). For the characterization of neuronal response types, we calculated the correlation of neural activity traces with each of 52 regressors formed to identify neurons primarily coding for different directions, eye muscles, eye position or OKR slow-phase eye velocity (Supplemental Fig. 1a-a’). We found that eye-correlated neurons are virtually always active during clockwise or counter-clockwise binocular stimulation (2380 out of 2508 neurons, from 15 larvae with each recording depth sampled 8-fold). They only differ from each other with regard to the extent of recruitment during monocular eye movements (Fig. 4c, Supplemental Fig. 1e).

**Figure 2:**
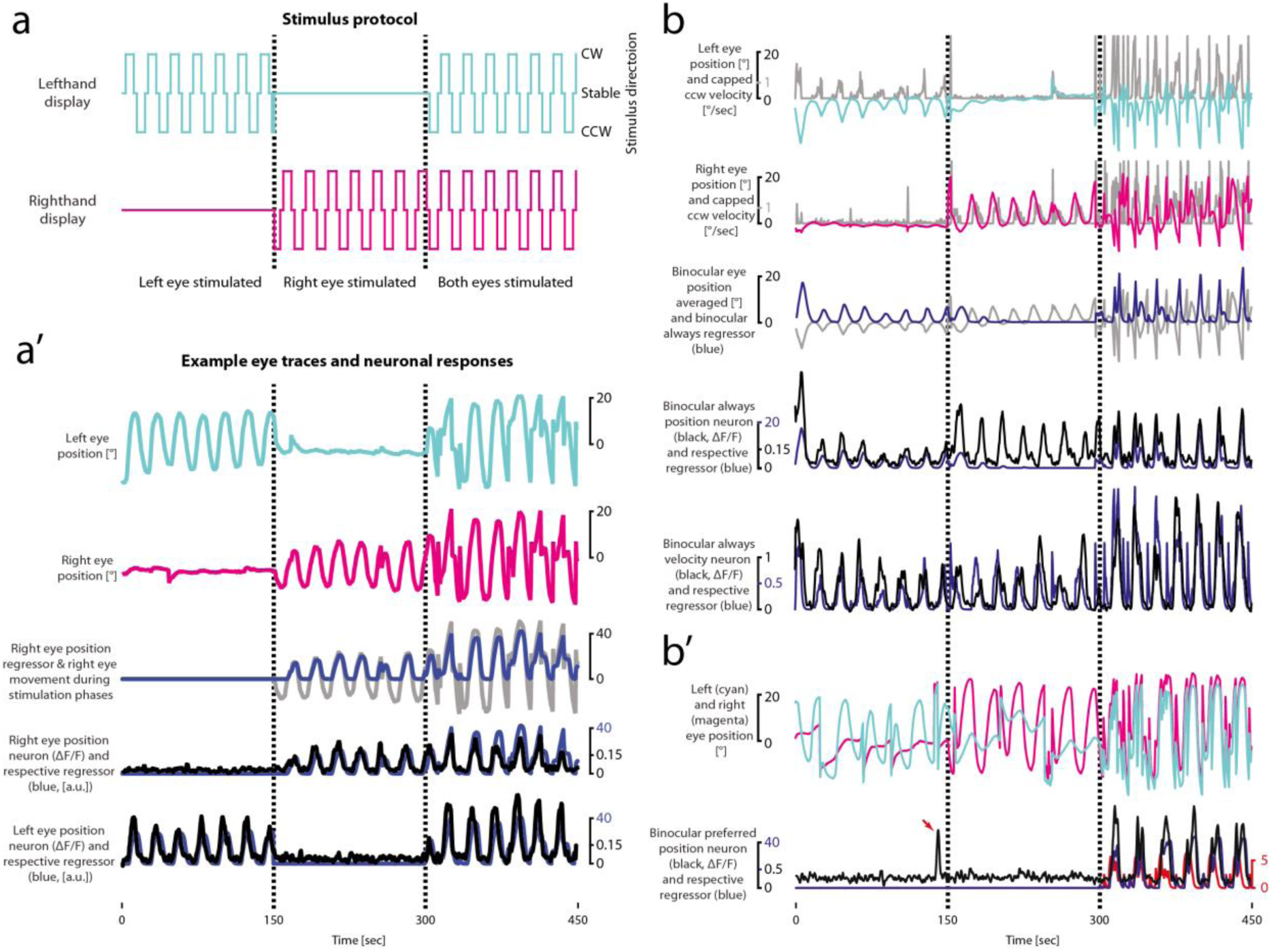
Experimental strategy to assess binocular coordination. **a:** Stimulus protocol for data shown in a’. Lines indicate direction in which the stimulus is moving. Dashed lines separate stimulus phases. **a’:** Example eye traces (right eye: magenta, left eye: cyan) and corresponding neuronal calcium responses (black, ΔF/F) with monocular coding. The respective highest scoring regressor [Monocular right eye, rightward eye position (r3); monocular left eye, rightward eye position (r7)] is shown in blue. The grey line shows right eye position from which r3 was derived. **b**: Example calcium responses of binocular neurons. Left (cyan) and right (magenta) eye traces with capped counter-clockwise eye velocity (grey, upper two plots) and averaged eye position (grey, third plot from the top) of which regressors r14 (binocular always leftward velocity) and r18 (binocular always leftward position) were derived. Black lines show ΔF/F for a binocular always (BA) position (P) and a BA velocity (V) neuron with the corresponding highest scoring regressor in blue. Note that the eye position for the right eye was mostly shifted towards the right side which resulted in almost no activity for R18 in the middle phase, although the regressor still classified the neurons correctly. **b’**: Example binocular preferred (BP) position neuron with respective eye trace; note the binocular event during the left eye stimulation and the corresponding activity (red arrow). The blue trace shows the respective regressor (binocular preferred, rightward position, r1), the red trace the corresponding velocity regressor (binocular preferred, rightward velocity, r9).

We identified four primary response types in our hindbrain data: two monocular (M) types with activity for either the left or the right eye (LE, RE), which were also active during the binocular stimulus phase (MLE, MRE, Fig. 2a’, Fig. 3a-b, Supplemental Fig. 3a-b), and two binocular response types. The binocular response types (types BA and BP, Fig. 2b-b’ and Fig. 3c-d) were either always active (‘binocular always’, BA, Fig. 2b), or showed a preference towards binocular eye movements (‘binocular preferred’, BP, Fig. 2b’).

**Figure 3:**
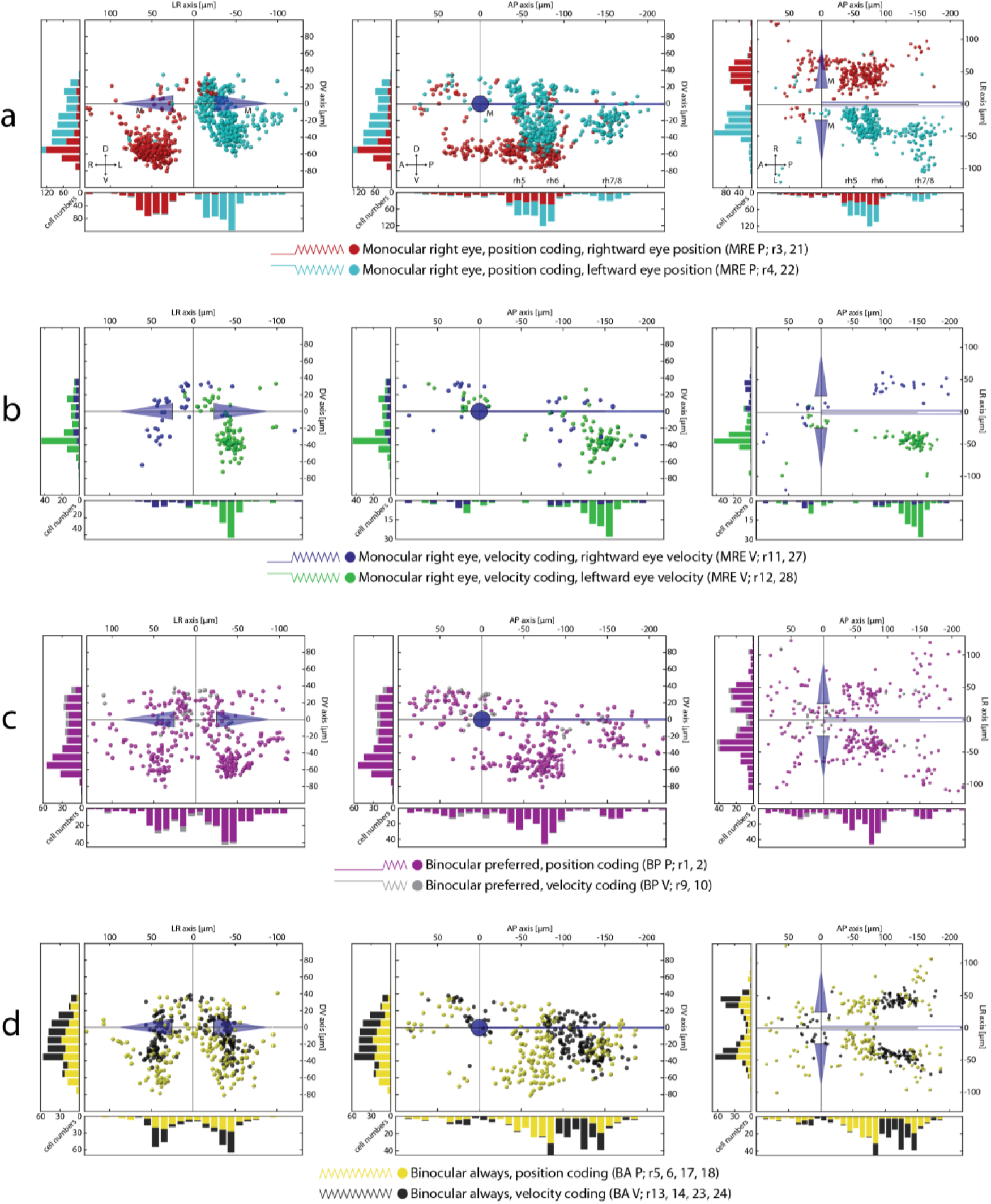
Monocular and binocular cell maps. **a-d:** Transversal, sagittal and dorsal views for MRE and binocular neurons in the hindbrain (see Suppl. Fig. 3a-b for mirror-symmetric MLE neurons). A: anterior; BA: binocular always; BP: binocular preferred; D: dorsal; L: left; M: Mauthner cells; MRE: monocular right eye; P: position/posterior; R: right; r: regressor; rh 5-8: rhombomeres 5-8; V: ventral/velocity; each coloured ball represents one neuron identified in one fish.

Since the motor range for eye movements during the binocular stimulation phase was mostly larger than during the monocular phases, we excluded all neurons that did not reach their firing threshold during the monocular phase (Supplemental Fig. 2).

98 % of eye movement correlated neurons, caudal to the Mauthner cells, responded in an ipsiversive manner (2110 vs. 37), though this hemispheric restriction was less prominent rostral to the Mauthner cells (65%, 228 vs. 133). Eye movement correlated neurons on the right side of the hindbrain are increasingly active during rightward eye positions (of the left and/or right eye) and vice versa.

### Monocular neurons

Monocular position encoding neurons are primarily located in rhombomeres 5 and 6 – based on mapping performed using the HGj4a line [31] –, forming two distinct columns in each rhombomere (Fig. 3a; Supplemental Fig. 3a). A second cluster can be seen around 150 µm caudal to the Mauthner cells and 40 µm lateral to the medial longitudinal fasciculus (MLF). This region in rhombomere 7/8 partially overlaps with the areas previously described as the OI in zebrafish [12]–[14], extending caudal-ventrally into the inferior olive (IO) which we find is mostly monocular encoding. The putative OI region contains a high number of position neurons encoding the contralateral eye and only few neurons encoding the ipsilateral eye.

Within our imaged brain volume containing rhombomeres 5 and 6 position neurons coding for the ipsilateral eye span only a narrow band 30 to 70 µm ventral to the MLF. Based upon the wiring diagram (Fig. 1a) and their response profile (ipsilateral, ipsiversive, position coding), these neurons located in the ABN correspond to motoneurons innervating the lateral rectus. Internuclear neurons carrying the information used to innervate the medial rectus should be located on the contralateral side and respond to contraversive positions. Such putative INNs are abundant and located more medially and dorsally than motoneurons, spanning a wider range from 60 µm ventral to around 30 µm dorsal to the MLF. These two clusters of putative moto- and INNs in the ABN are mirror-symmetrical between monocular left and right eye encoding neurons (Fig. 4a). Monocular contralateral encoding neurons showed a volume with fewer neurons 10 to 30 µm ventral to the MLF rotated roughly by 20 degrees along the RC-axis, separating them into two groups (black arrows/inset Fig. 4a).

**Figure 4:**
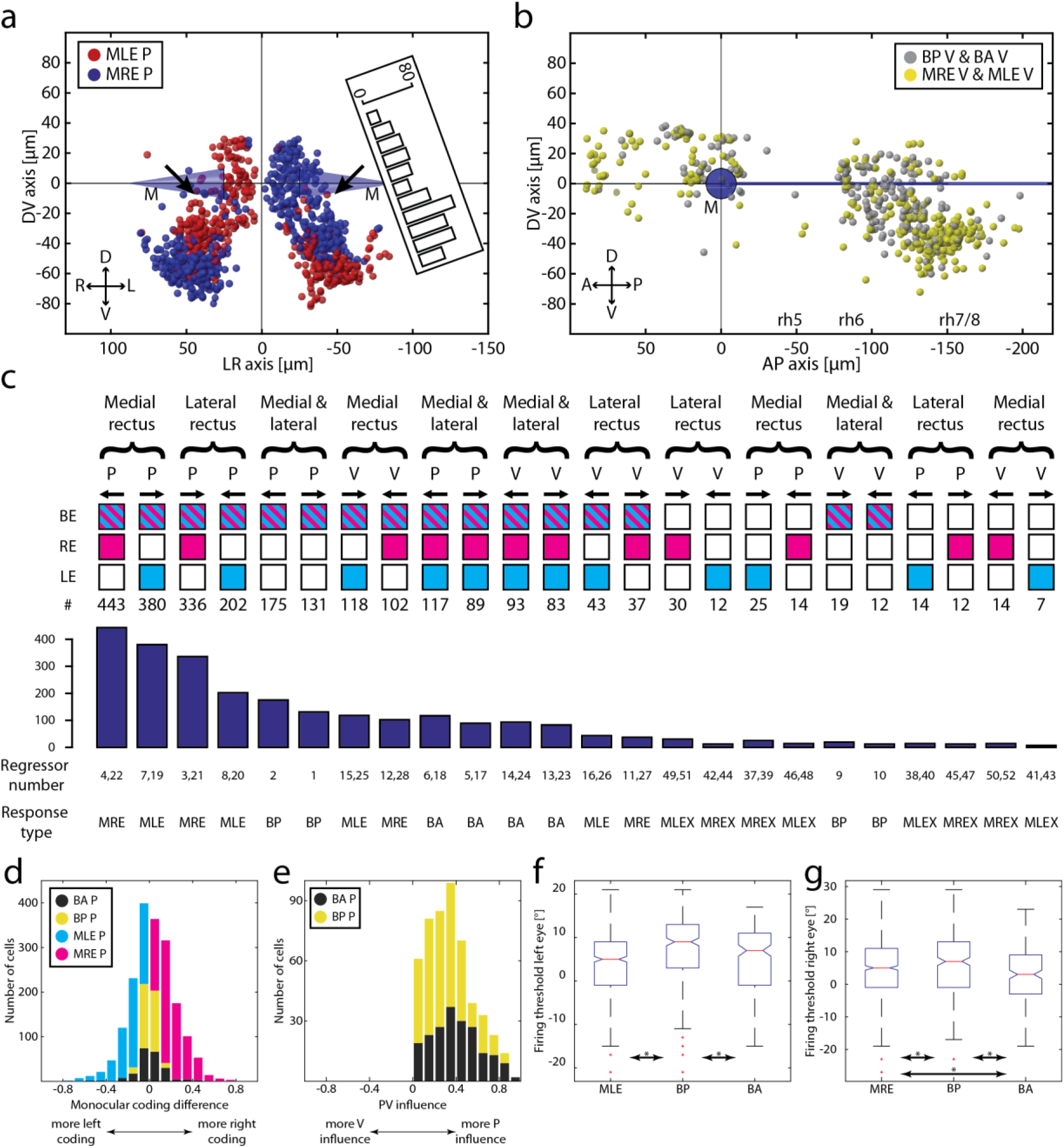
Monocular/binocular synopsis. **a:** Transversal projection of monocular coding neurons within rh5/6 (ABN). D: dorsal; L: left; M: Mauthner cells; MLE: monocular left eye; MRE: monocular right eye; P: position; R: right; V: ventral. Black arrows indicate position of a faint gap between the ventral and dorsal neurons. Inset shows the numbers of neurons for the left hemisphere along the D-V axis rotated by 20°. **b:** Monocular and binocular velocity encoding neurons. A: anterior; BA: binocular always; BP: binocular preferred; P: posterior; rh 5-8: rhombomere 5-8; **c:** Number of neurons found for each response type sorted pairwise according to the affected muscle(s). BA: binocular always; BP: binocular preferred; MLE: monocular left eye; MLEX: monocular left eye exclusive; MRE: monocular right eye; MREX: monocular right eye exclusive P: position; V: velocity; **d:** Monocular coding differences for all four main response types for position coding neurons. Index running from −1 (exclusively coding for left eye) to +1 (right eye); **e:** PV influence for BA P and BP P neurons. Index running from −1 (exclusive velocity influence) to +1 (exclusive position influence); **f-g:** Left and right eye firing thresholds acquired during the firing threshold analysis pooled in ON direction.

Monocular slow phase eye velocity neurons are mainly located ventrally to the MLF in rh7/8 and code for the contralateral eye. They are clustered slightly ventro-rostrally to the putative OI position neurons with some overlap between both clusters. As is the case for the monocular position neurons, the rh7/8 region contains only few monocular velocity coding for the ipsilateral eye. Rostral to these identified velocity neurons, some sparse, ungrouped neurons are located in both hemispheres, extending to the caudal end of rh6 (Fig. 3b; Supplemental Fig. 3b).

Monocular neurons preferentially active during one monocular stimulation phase and silent during binocular movements (monocular exclusive) were heavily underrepresented for both position and velocity (159 of 2508, Supplemental Fig. 4). Neurons exclusively active during both monocular stimulation phases were virtually absent (Supplemental Fig. 1a & 1e).

### Binocular neurons

We identified binocular neurons that were always active (BA) or were preferentially active during the binocular eye movements (binocular preferred, BP). The vast majority of BP neurons encode eye position, not velocity (Fig. 3c). They overlap with monocular position coding neurons in rhombomere 7/8, but their centre of mass is biased to a more lateral position. The rightward and leftward tuned BP neurons are distributed in the right and left hemispheres, respectively, as expected from the ipsiversive coding scheme. In the ABN, BP neurons were clustered more ventrally with more neurons in the left hemisphere than in the right (100 vs. 144; caudal to the Mauthner cells).

Binocular position neurons active regardless of stimulated eye or stimulus phase (BA) are homogeneously distributed in the ABN and putative OI (Fig. 3d), following the pattern of their monocular counterpart, and no lateralization across hemispheres was observed. However, those BA neurons that encode velocity form a narrow band (Fig. 3d, right panel) spanning from the dorsal end of rh6 (within our imaged region) to the location of monocular velocity coding neurons in rh7/8 and are absent from the remaining ABN and caudal rh7/8 region.

While BA neurons responded during all stimulus phases, their responses during monocular stimulus phases were typically smaller than those during binocular stimulus phases, which can likely be attributed to the smaller explored motor range during monocular stimulation (for an assessment of response type classification see discussion in the Methods section, Supplemental Fig. 3d).

While monocular and binocular position neurons shared the same anatomical locations in the zebrafish hindbrain, an anatomical response type gradient existed for velocity neurons caudal to rh6 (Fig. 4b): binocular velocity neurons are located more rostro-dorsally while monocular velocity neurons formed a cluster in the ventral part of rh7/8.

Having identified the four primary response types, we next sorted all occurring response types according to the number of identified neurons for each response type and grouped them according to the encoded eye direction (CW, CCW), controlled eye muscles (lateral rectus, medial rectus, or both) and kinematic parameter (eye position or OKR slow-phase velocity). This analysis (Fig. 4c) reveals that i) position neurons are more frequent in the hindbrain than slow-phase eye velocity neurons (1938 position vs. 570 velocity), ii) more monocular neurons coding for the medial rectus exist than monocular neurons coding for the lateral rectus eye muscle (1043 medial vs. 618 lateral), and iii) using our stimulus protocol we found more neurons coding for the position of the right eye than for the left eye position (779 right vs. 582 left; this might have been caused by a history dependence, as in 90 % of the recordings the left eye was monocularly stimulated before the right eye). For all mono- and binocular regressors we find neurons dorsal to the MLF and rostral to the Mauthner cells which show an intermingled anatomical distribution of ipsiversive and contraversive response types. This cluster corresponds to the caudal end of the previously described “hindbrain oscillator” [also termed ARTR, [3], [5], [6]].

To check how tightly the neurons are correlated to each specific eye, we calculated - for all four major groups - the difference in the correlation to the left and right eye (Supplemental methods). As expected, binocular neurons were located in the centre and had a unimodal distribution, while monocular neurons had a more bimodal distribution caused by the left and right coding population [Fig. 4d, Index running from −1 (more monocular left eye coding) to 1 (more monocular right eye coding)]. When comparing the velocity influence of BA (n=206) and BP (n=306) position coding neurons (Supplemental methods, Fig. 4e) we found that both groups showed similar velocity-position distributions, with BA position neurons having a slightly stronger position component than BP position neurons (two-sided Wilcoxon rank sum test, p=5.7*10^-7^, Index running from −1 (Velocity) to 1 (Position)). The firing thresholds (from the firing threshold analysis, Supplemental Fig. 2) of BP position neurons were shifted towards the ON direction compared to BA and monocular position neurons and, for the right eye, BA neurons showed significantly earlier thresholds than MRE neurons (Fig. 4f-g). While the response type classification used in this study (Fig. 1-3, Fig. 4a-c) is instructive for understanding the processing repertoire of the oculomotor hindbrain, the results presented in Fig. 4d-g show that the responsivity of oculomotor neurons forms gradients within the parameter space spanned by the regressors used for our response type classification. For example, the BP Position classification could be affected by velocity components and a larger dynamic range of eye positions during the binocular stimulation phase, and furthermore some BP neurons were also active during the monocular stimulation phases, albeit at low activity levels preventing their classification as BA or monocular. Taken together, this suggests that BA and BP neurons might not be two distinctively separate groups but that they exist along a continuum, with the extreme cases being BA and BP.

### Differential encoding of velocity and position in individual neurons

Our first experiment was geared towards identifying monocular versus binocular tuning. We also classified neurons as either mainly position or mainly velocity encoding (Fig. 3) in this experiment, although intermediate ‘multi-dimensional’ responsivity likely occurs as well. ABN neurons should receive slow-phase velocity signals during optokinetic stimulation, e.g. via the pretectum/accessory optic system, vestibular nuclei and the OI [Fig. 1a’; [20], [32]–[35]] since a muscle force step is needed to overcome the dampened, viscous kinetics of the oculomotor plant [36], [37]. In order to investigate the differential coding of oculomotor neurons and to visualize the anatomical distribution of position and velocity coding within rhombomeres 7/8, we developed a binocular closed-loop stimulation protocol to disentangle eye position from eye velocity correlations by eliciting different eye velocities at different eye positions (Fig. 5a-a’’, Methods). This allowed us to consistently evoke combinations of eye position and velocity which would only occur sporadically during optokinetic responses to fixed stimulus sequences. At the same time the stimulus protocol minimized the occurrence of fast phase eye movements (saccades) in order to improve our ability to relate neuronal activity to slow phase behaviour in this correlative experiment; i.e. the experiment was not designed to identify or characterize the burst system responsible for generating saccades [3], [38]. From the whole recording we constructed two-dimensional tuning curves covering the activity for almost all different eye position and slow phase eye velocity combinations within a certain range (eye position: −15° to +15°, eye velocity: −7 to +7 degrees/sec, Fig. 5b-d, Supplemental Fig. 5a-c). Using this protocol we analysed 889 neurons, which exhibited different combinations of eye position and slow-phase eye velocity tuning. To classify the differences in position and velocity coding for each neurons we calculated a Postion-Velocity index (PV_Index_) based on the correlation of the neuronal response to behavioural regressors (see Methods). This index runs from −1 (pure velocity coding) to +1 (pure position coding). Both neurons tuned exclusively to position (neurons 1) or velocity (neuron 3) exist, as well as intermediate cases (neuron 2, Fig. 5b-d). For neurons with an intermingled position and velocity component (-0.5 < PV_Index_ < 0.5) the preferred direction was almost always the same for position and velocity (94%, 440/470).

**Figure 5:**
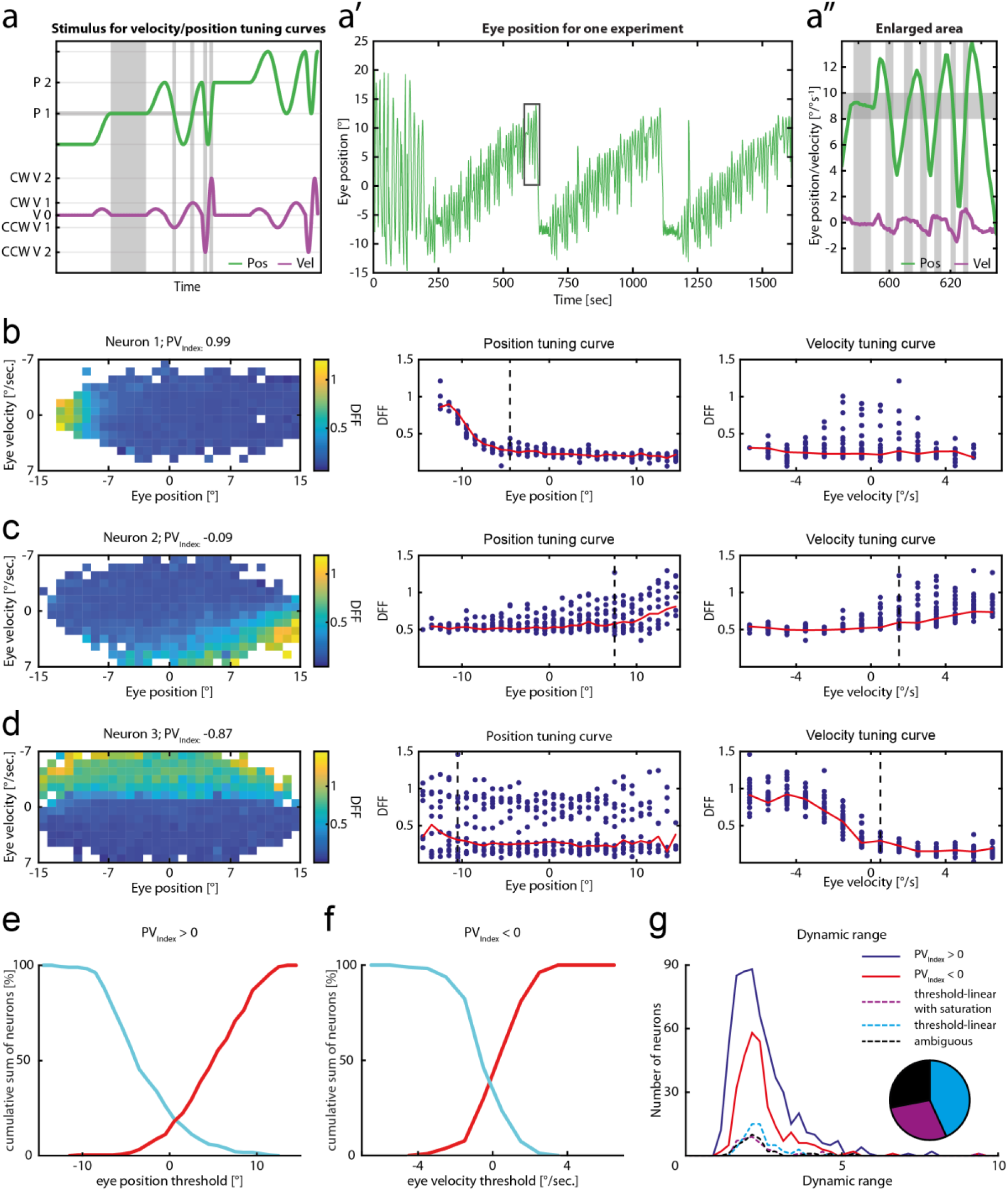
Neuronal tuning for eye velocity and position. **a:** Schematic of the closed loop velocity/position stimulus for highlighted eye position (P1) at different slow-phase eye velocities (CCW V2, CCW V1, V0, CW V1, CW V2). Only two velocity steps are depicted for illustration purposes. Grey shaded rectangles show one eye position bin and different velocities for that bin. CCW: counter-clockwise; CW: clockwise; P: position; V: velocity **a’:** Example binocular eye trace for one recording. **a’’**: Highlighted area from a’. Grey boxes as in a. **b-d:** Left panel: Tuning curves showing DFF colour coded for averaged eye position-velocity bins. Middle panel: Position tuning curve. Red line shows averaged DFF between ± 0.5 °/sec eye velocity, blue dots for every other eye velocity bin (as in left panel). A black dashed line shows the firing threshold, if identified. Right panel: same as for the middle panel, but for eye velocity. Red line shows averaged DFF between ± 2° eye position. **e**: Cumulative position threshold plot for position coding neurons (PV_Index_ > 0) pooled in ON direction to the right (red, n=250) and left (cyan, n=283). **f**: Cumulative velocity threshold plot for velocity coding neurons (PV_Index_ < 0) pooled in ON direction to the right (red, n=104) and left (cyan, n=175). **g**: Dynamic range of fluorescence for position and velocity coding neurons (PV_Index_ > 0, PV_Index_ < 0 respectively) and for neuron with a very strong velocity coding (PV_Index_ < −0.5, dashed lines) separated by their response profile. Pie chart showing the relative numbers for strong velocity coding neurons (w/ saturation: 29 % (40/139), w/o saturation: 43% (60/139), ambiguous: 28 % (39/139)).

### Firing thresholds of positon neurons are distributed across the dynamic range of eye positions while velocity neurons mainly get activated at velocities close to 0°/sec

To quantify the neuronal recruitment we used the two-dimensional tuning curves and analysed the position and velocity firing thresholds in the position and velocity planes intersecting with the origin. This procedure results in one-dimensional eye position tuning curves around eye velocities of 0°/s (black and red line in Fig. 5b-d middle panel) and eye velocity tuning curves around eye positions of 0° (right panel) for the same neurons. For position encoding neurons (PV_Index_ > 0, n=533 neurons with identified position threshold) we find that the eye position thresholds were distributed across the full motor range (roughly - 10° to +10°, Fig. 5e). Leftward and rightward eye position encoding neurons had slightly different eye position thresholds in our dataset [Wilcoxon rank sum p= 0.000016, median for rightward coding neurons pooled on ON direction (n=250): 5.5, leftward 4.5 (n=283)]. For the velocity encoding neurons (PV_Index_ < 0, n=279) the activation thresholds for velocity mostly span a small range mostly between ± 2°/s, so that the calcium signals started to increase at eye velocities close to 0°/sec. No difference was observed between velocity neurons coding for leftward vs. rightward velocities (Fig. 5f, Wilcoxon rank sum p=0.24; rightward n=104, leftward n=175). For the majority of velocity neurons, the velocity tuning curve did not cross the velocity of 0 °/s, i.e. the neurons were only active for either positive or negative velocities. Also the strongest fluorescence increases were usually observed after crossing a velocity of 0°/s. However, the true firing thresholds may start further into the OFF direction (≤0°/s) as we likely couldn’t reliably detect single action potentials using GCaMP6f in our preparation [39].

Visual inspection of all strong velocity neurons (PV_Index_ < −0.5) revealed that some of the identified velocity neurons showed firing saturation at higher velocities (29 %; 40 of 139; Fig. 5g). Calcium indicator saturation, which occurs at high calcium concentrations ([Ca]^2+^>>K_d_), is unlikely to account for the observed fluorescence saturation, since the dynamic range of fluorescence values (F_Max_/F_Min_) was (i) much smaller (∼2.5) than the published range of the GCaMP6f indicator (51.7) [39], and (ii) similar for non-saturating position neurons and saturating velocity neurons (Fig. 5g).

For neither of the two velocity tuning types (saturating vs. non-saturating) a clear anatomical clustering is visible (Supplemental Fig. 6) and we therefore merged the corresponding neurons into one group (potentially the non-saturating neurons could still saturate at higher eye velocities not reached in our experimental protocol).

**Figure 6:**
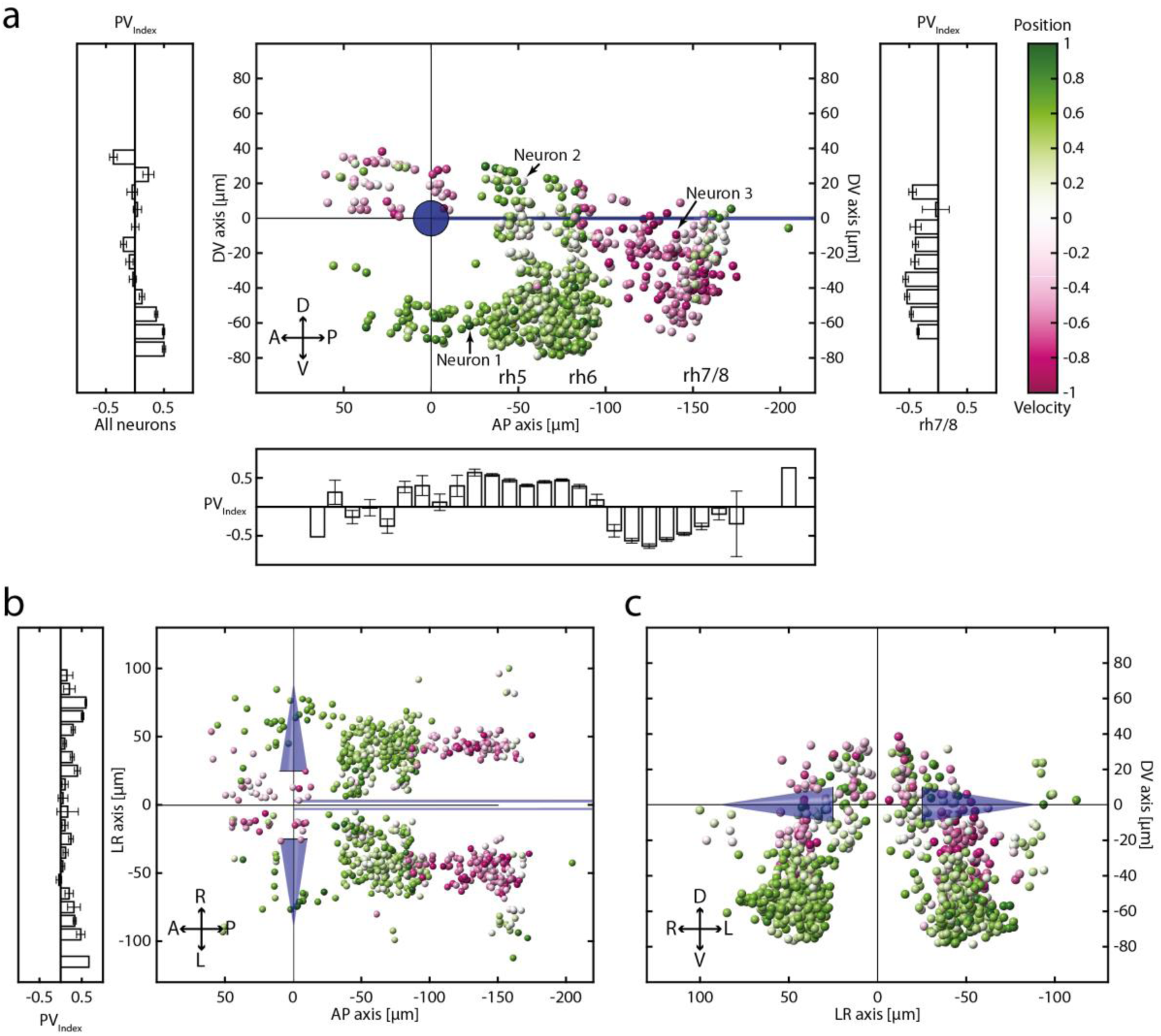
PV_Index_ distribution and spatial location of identified neurons. **a-d**: Sagittal (a), dorsal (b) and transversal (e) anatomical views of eye-correlated neurons color-coded for the PV_Index_. Histograms show the anatomical distribution of neurons along the appropriate axis for either all neurons or exclusively for rh7/8. Blue cones: Mauthner cells, blue line: MLF; A: anterior, D: dorsal; P: posterior; V: ventral; Error bars are SEM.

### No anatomical gradients of oculomotor tuning thresholds in the hindbrain

In order to investigate topographical arrangements of tuning thresholds in the hindbrain, we generated anatomical maps of firing thresholds for position (P_Thres_) and velocity (V_Thres_) for position neurons with an identified threshold (PV_Index_ > 0, n=533, Supplemental Fig. 7a) and for velocity neurons (PV_Index_ < 0, n=279, Supplemental Fig. 7b). Position thresholds do not appear to be anatomically grouped and no clear anatomical gradient within any of the neuronal clusters could be identified (Kruskal-Wallis test for position threshold differences p=0.07; rh5: 214; rh6: 249; rh7/8: 27). We investigated whether MNs (based on anatomical location) are distributed topographically according to position firing threshold, but were unable to identify a significant gradient [Kruskal-Wallis p=0.22, Supplemental Fig. 7a].

Eye velocity thresholds (V_Thres_) also did not show any spatial clustering and no gradient could be observed within the hindbrain. No statistical difference was observed (Kruskal-Wallis p=0.79; rh5: 11; rh6: 10; rh7/8: 184).

### Neurons in rhombomere 7/8 exhibit a velocity-to-position gradient

The anatomical clusters of position and velocity coding neurons that we identified using the PV_Index_ are generally in agreement with those obtained from the separate experiment described above (compare Fig. 6a-c to Fig. 3 & Supplemental Fig. 3). Neurons in the ABN (rh5/rh6) displayed an average PV_Index_ of 0.44 (±0.23 STD; n=521) indicating position tuning with some minor velocity sensitivity. Within the ABN, the velocity component was strongest around the previously described gap (see Fig. 4a, black arrows) in-between two clusters of neurons 20 µm ventral to the MLF. The velocity neurons identified using the velocity-position stimulus reside in the ventral part of rh7/8 and extend into the area caudal to rh6, overlapping with the volumes containing the BA, MLE and MRE velocity neurons (Fig. 3b-d, Supplemental Fig. 3b). In the caudal part of rhombomeres 7/8 we find neurons with more position coding dependence than in the rostral part, especially laterally (Fig. 6a-c). Following the anterior-posterior and ventral-dorsal axes in the caudal hindbrain (rh7/8), our analysis therefore reveals a prominent PV_Index_ gradient, shifting from velocity towards an intermingled velocity/position tuning with neurons exhibiting a stronger position coding at the dorso-caudal end (Fig. 6a).

## Discussion

We investigated the binocular coordination, eye velocity and position sensitivities, as well as associated recruitment orders and anatomical distributions of oculomotor neurons in the zebrafish hindbrain.

We find four predominant response types, comprised of two monocular and two binocular types (Fig. 7). Monocular neurons consist of MNs, INNs, putative OI, VSM and IO neurons. We found that abducens INNs are mainly located dorsally to the MNs (Fig. 4) and together mainly code for eye position (Fig. 7b). In the caudally adjacent rhombomeres 7 and 8, oculomotor neurons mainly code for eye velocity and form a rostro-caudal velocity-to-position gradient. No clear segregation between velocity and position encoding neurons could be identified in this volume, suggesting that oculomotor integrator and the velocity storage mechanism merge smoothly at this developmental stage. A large fraction of neurons preferentially encode binocular eye movements showing that the recruitment of neurons depends on the executed behaviour (monocular or binocular OKR). Given the number of identified neurons, those coding monocularly for the lateral rectus in OI and VSM are almost absent (Fig. 4, Fig. 7c), which is discussed further below.

**Figure 7:**
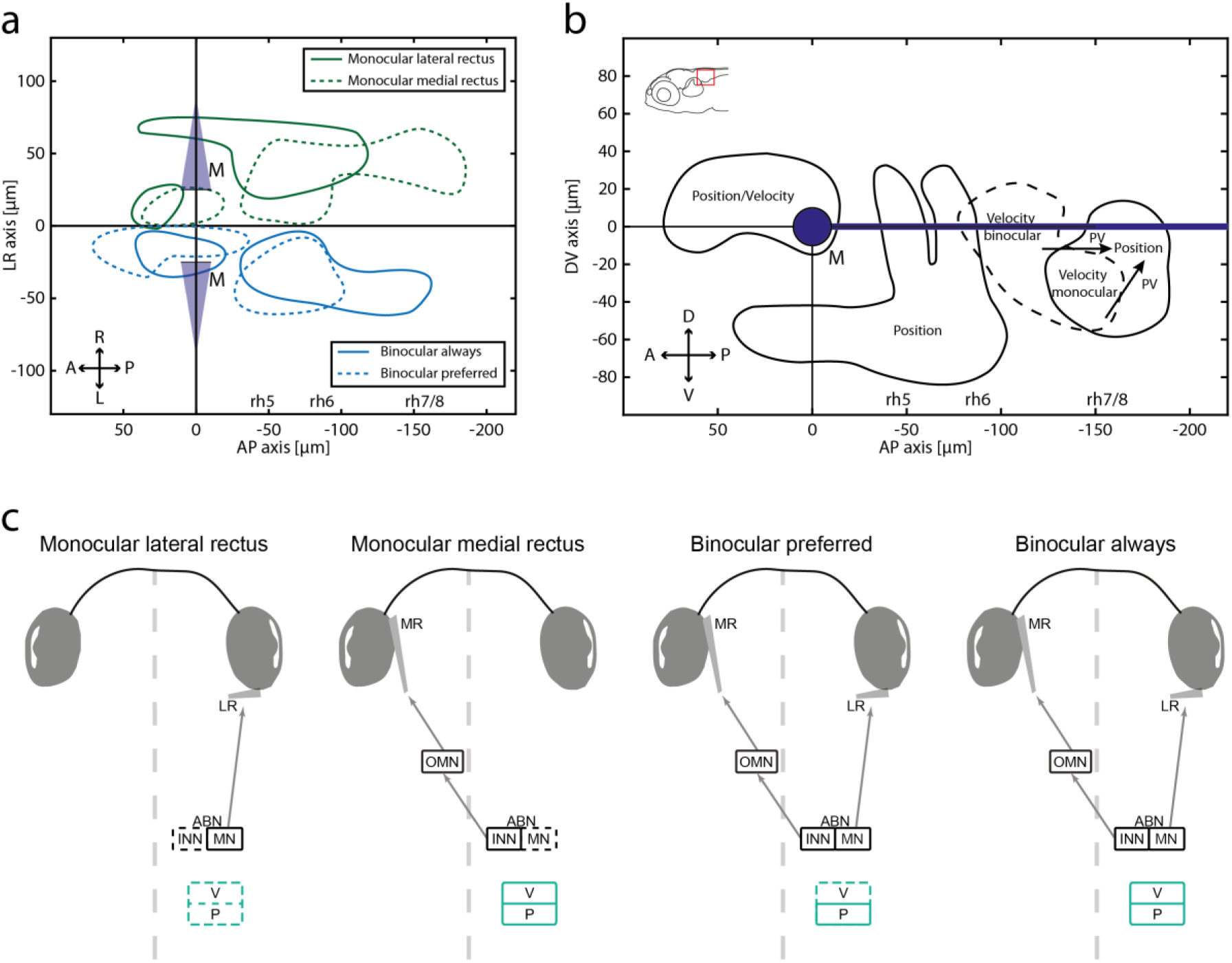
Summary for binocular coordination and PV encoding in the larval zebrafish hindbrain. **a:** Anatomical separation of monocular and binocular neurons in the dorsal view. For illustrative purposes, all monocular domains are depicted in the right hemisphere, and binocular domains in the left hemisphere (no difference across hemispheres was identified). A: anterior; L: left; M: Mauthner cells; P: posterior; R: right; rh5-8: rhombomere 5-8. **b:** Distinct clusters of eye movement coding neurons in the hindbrain (side view). Arrows indicating position-velocity shift in the OI. D: dorsal; V: ventral. **c:** Schematic illustrating each response type. Note the absence of slow-phase velocity neurons with preferred binocular (BP) encoding and the lack of monocular neurons for the temporal half of the ipsilateral eye outside of the nucleus abducens. Dashed lines represent “missing” neuronal clusters, i.e. only a small numbers of neurons were found for the respective eye movements.

### Anatomical organisation of MNs and INNs in the ABN

To reveal the anatomical volumes containing MNs and INNs in the ABN we made use of the fact that the lateral rectus eye muscle is innervated by ABN MNs and should increase its activity during ipsiversive (temporal/abducting) movements of the ipsilateral eye. We report the location of MNs to be limited to the ventral ABN, which is in line with transgenic marker lines for mnx1+ motoneurons [*vu504Tg*, [40]]. The INNs are located more dorsally with only a small intermingled zone between the MNs and INNs. This is in line with data from goldfish where ventral MNs and more dorsal INNs form 4 separate clusters with 2 of them being adjacent and - to some extent - intermingled with each other [41], [42].

In our data we see a faint gap (20 µm ventral to the Mauthner cells) running along a dorso-lateral to medio-ventral axis in the cluster of putative INNs, which separates them into two groups (black arrows Fig. 4a). While the dorsal and the ventral domain both carry mainly the same information (contralateral eye encoding, ipsiversive eye positions), the dorsal group is in close proximity to a group of neurons recently investigated and identified as the medial vestibular nucleus (MVN) by D. Schoppik and colleagues [[43], which has been registered in the z-brain atlas using the *Tg(-6.7FRhcrtR:gal4VP16)* line [44]]. However, our dorsal group of neurons covers a larger volume and extends more medially than the annotated MVN in the z-brain atlas and mainly codes for eye position, not slow-phase velocity. It is nonetheless possible that the dorsal group partially corresponds to the MVN.

Very ventrally we find a group of neurons extending rostrally from the pool of rh5 MNs coding for eye position monocularly and binocularly (Fig. 7b, [40 to -40 µm on AP axis, −60 µm on DV axis]). As they are not located in the ABN nor labelled in a line specifically labelling MNs (*vu504Tg*), these neurons likely do not project to the extraocular muscles and instead might carry efference copy signals.

### Anatomical organisation of the caudal hindbrain (rhombomeres 7/8)

Neurons at the ventral-caudal end of the hindbrain were located very close to the floor plate of the brain, and overlapped with the anatomical location of the inferior olive [45], as were neurons more than 70 µm lateral from the MLF in the caudal hindbrain. We did not see a clear anatomical-functional segregation of eye-movement correlated putative OI and inferior olive neurons. Our results and the previous studies suggest that within our cluster of oculomotor neurons in rh7/8, the medial-rostral, as well as the medial-caudal-dorsal neurons correspond to the OI, while the ventral-caudal neurons correspond to the inferior olive [compare Fig. 5g-j in [46], Fig. 2 in [13]]. Comparing the medio-lateral extent of our putative OI neurons we do not find neurons closely located to the midline as shown in other studies [12]–[14], [46]. As these medially located neurons were reported to be located more dorsally, our recordings might have missed such neurons in dorso-caudal regions. However, in a recent EM study medially located neurons have been found exclusively at the rostral end of rh7 [boundary to rh6, Fig. 1d and Supplemental Fig. 3 in [14]], an area which we extensively imaged and which contains many velocity sensitive neurons (rh7) as well as position sensitive neurons in rh6 (ABN/MVN).

The axonal projection patterns of our reported functional neuron types remain to be identified. The majority of our OI neurons are located ventral to the MLF, likely overlapping with the glutamatergic stripes 1 & 2 [Fig. 2a in [13]] and the GABAergic stripe S2, which contain both ipsilaterally and contralaterally projecting axons.

### Lack of monocular coding for the lateral rectus muscle in the caudal hindbrain

We show that monocular neurons in rhombomeres 7/8 almost exclusively encode the contralateral eye in larval zebrafish. In monkeys it was reported that 50 % of monocular burst-tonic neurons in the nucleus prepositus hypoglossi (NPH) and medial vestibular nucleus (MVN, mammalian equivalents to the OI) code for the ipsi- or contralateral eye during disjunctive fixation/saccades [47], while another study reports “most” (*sic*) of monocular NPH neurons to be related to the ipsilateral eye [48]. Data from goldfish also shows that only 4% of neurons in Area I (equivalent to OI) code for the contralateral eye and 57 % for the ipsilateral eye during monocular stimulation [49].

Therefore, a lack of monocular coding for one extraocular eye muscle can be observed in the oculomotor integrator. While Debowy and colleagues [49] find almost no monocular integrator neurons coding for the nasal part (medial rectus) of the contralateral eye in goldfish, in the present study we are missing monocular neurons encoding the temporal hemisphere (lateral rectus) of the ipsilateral eye (Fig. 7c). This part would only be encoded in the binocular context.

### A mixed, but task-specific monocular-binocular code

Almost all neurons described in this study were active during conjugate eye movements. According to Hering’s hypothesis monocular eye movements are not effected by monocular signals, but by the summation of binocular signals, which oppose each other in one eye and summate in the other eye, thereby effecting monocular eye movements by means of binocular conjugacy and vergence commands. While we did find BA neurons (whose response profiles are in line with conjugacy commands), the (almost complete) lack of neurons coding for vergence (which would be active only during disconjugate/monocular eye movements in our experiments) is in disagreement with Hering’s hypothesis. On the other hand, functional neuron types tuned to a single eye are abundant in the zebrafish hindbrain. These neurons are active regardless of whether the eye movement was monocular or conjugate and their existence conforms to Helmholtz’ hypothesis.

The functional structure of the zebrafish ABN shows that recruitment of neuronal pools depends on the executed OKR behaviour. The BP pool is preferentially activated during conjugate eye movements and less active during monocular eye movements. The anatomical location of the dominant cluster of BP neurons in the ventral part of the zebrafish ABN, intermingled with monocular coding neurons, suggests that many of these BP neurons are indeed MNs. The fact that ABN motoneurons differ in their eye preference and also encode binocular information has previously been shown in monkeys by W. M. King and colleagues [[48], [50]–[52], discussed in [53]]. The functional classification (monocular or binocular encoding) thus does not necessarily correspond to the connected extraocular eye muscle, as ABN motoneurons connect exclusively to the LR muscle of the ipsilateral eye. Our finding represents a deviation from a strict final common pathway: neurons coding for the same eye in different behavioural contexts (binocular vs. monocular OKR) are differentially recruited in these two contexts. Furthermore, if an extraocular motoneuron gets recruited only in certain behavioural contexts (e.g. conjugate eye movements), the lack of motoneuron activity for the innervated eye (e.g. during monocular eye movement) must be compensated by other neurons or elsewhere in the system [54]–[56] to maintain the eye position. Future studies are needed to reveal how the oculomotor system reconciles this apparent paradox, and the small number of cells involved in the larval zebrafish could facilitate corresponding experiments.

### Recruitment orders for eye position and eye velocity

The analysis of one-dimensional tuning curves for eye velocity revealed that velocity encoding neurons in the zebrafish hindbrain each increase their firing for one out of the two directions tested, but are not strictly direction-selective: a minority of neurons already start firing during slow-phase eye movements into the non-preferred direction. This feature of eye velocity tuning has previously been observed in individual neurons of the goldfish Area II as well (cf. Fig. 7b in [23]). However, activations for non-preferred directions were mostly of small magnitude in our data and it remains possible that recording noise or sampling errors affected the identified velocity thresholds. Due to the above described saturation of velocity signals, a fraction of velocity neurons exclusively encode information for very slow eye velocities, which might enable more precise control of eye velocity in the velocity regime close to 0°/s. The eye position firing thresholds of position neurons, however, distribute across the dynamical range of eye positions, which is in agreement with previous reports on the recruitment order in the ABN and OI of other species [18]–[20], [57]–[59]. Our analysis of tuning thresholds did not reveal any anatomical gradients for these eye position and velocity thresholds. This includes the MNs located in the ABN (Supplemental Fig. 7a) for which a soma size gradient has been reported recently [60].

### The existing correlations to retinal slip signals remain to be investigated

In order to generate many and quickly changing eye movements within the limited recording time of our experiments, we chose to use relatively high stimulus velocities. This caused low optokinetic gains [24] and considerable error signals resulting from the remaining retinal slip during slow-phase eye movements. Next to the eye velocity correlations which we describe in this study, these slip signals correlate with the activity of velocity neurons as well. We checked the full dataset of the velocity/position experiment and found that only 4 out of 635 neurons showed a better correlation to a retinal slip signal than to eye position or velocity (correlation analysis, data not shown).

### Persistent activity generation likely relies on the observed velocity-to-position gradient in the caudal hindbrain

Our analysis of differential position versus velocity encoding (PV_Index_) revealed dominance of position coding in the ABN (rh 5/6) and an anatomical velocity-to-position gradient of oculomotor neurons in rhombomere 7/8, which have stronger velocity weights in the rostral part of rhombomere 7/8 and stronger position weights in the caudal part.

The rostral part of our identified velocity coding neurons (in rh7) likely corresponds to the velocity storage mechanism [Area II in fish, [21], [22]], which is rostrally adjacent to the OI (Area I) in goldfish. While in adult goldfish a clear functional separation of Areas I and II has been reported, in the larval zebrafish, the velocity and position encoding in rh7/8 appears to form a gradient, making it difficult to draw a border between the velocity storage mechanism and the OI. While the velocity storage mechanism is still maturing in 5 dpf old larval zebrafish (it only stores the velocity for one or two seconds as measured using the optokinetic after-nystagmus [[25], and own observations]), the hindbrain already contains a high number of velocity coding neurons.

Our data suggests that the velocity-to-position gradient extends well into the anatomical region of the OI and does not reach exclusive position sensitivity. Therefore the OI appears to perform only a partial integration (at this developmental stage), where the velocity signals are integrated into an intermediate velocity-position state [61], [62]. This gradient is in agreement with a previous publication which identified a change of persistence times in the OI along the rostral-caudal and dorsal-ventral axis [12]. These results suggest that integration is achieved by a feed-forward organisation of neurons, which gradually change in their position/velocity coding and persistence time. While partial integration can theoretically explain the heterogeneity and spatial gradients of time constants within the integrator some contradictions to integrator models still remain [63].

It has been previously reported that the activity of the zebrafish OI encodes two separate parameters [64]: while the amplitude of OI neuron activity represents eye position, the spatial pattern of persistent firing represents the context of how the eyes reached that position. If eye positions were reached during optokinetic behaviour, the rostral neurons of the OI showed more persistent activity, while during spontaneous saccadic movement the spatial pattern was reversed. Our results show that in parallel to the previously reported context-dependent anatomical gradient, slow-phase eye velocity is encoded in a similar gradient as well, such that (based on their anatomical rh7/8 location) neurons recruited during OKR are likely to also have a higher velocity sensitivity.

## Conclusion

Our findings characterize the functional layout of the oculomotor hindbrain in zebrafish. They reveal the functional oculomotor architecture regarding a set of key parameters (monocular/binocular encoding, position/velocity encoding, tuning curve/firing thresholds, anatomy) useful for future investigations into mechanisms underlying persistent activity and sensorimotor transformations. We provide evidence for a mixed but task-specific binocular code and suggest that generation of persistent activity is organized along the rostro-caudal axis in the larval hindbrain.

## Material & Methods

### Fish husbandry

Zebrafish (*danio rerio*) expressing GCaMP6f were used in the experiments [*Tg(ubi:nls-GCaMP6f)m1300*; Additional file 1]. Larvae were raised in a 14/10 h day/night cycle incubator at 29 °C in E3 solution containing methylene blue. Fish were kept in a TL/N [nacre; [65]] background, imaged larvae were *nacre -/-*.

### Transgenesis

The *Tg(ubi:nls-GCaMP6f)m1300* line was created using the Tol2 transposon system [66] and Gateway cloning (Invitrogen, 12537-023, Version D). Briefly, an attB1 primer (GGGGACAAGTTTGTACAAAAAAGCAG*GCTACC***ATGGCTCCAAAGAAGAAGCGTAAGGTA**TGGGTTCTCATCATCATCATC) including Kozak [67] and nls [68] sequences was used to amplify GCaMP6f [[39], Addgene plasmid #40755 pGP-CMV-GCaMP6f]; the ubi promoter [3.5 kb, [69], Addgene plasmid #27320] was inserted into the pENTR5’ plasmid. pENTR5’ (ubi), pME (nls-GCaMP6f) and pENTR3’ (polyA) sequences were then cloned into the pDestTol2pA2 plasmid via an LR reaction. 25 ng/µl plasmid DNA and 50 ng/µl Tol2 *transposase* mRNA were co-injected into single cell stage embryos (*nacre +/-*). F2 or fish of later generations were used for data acquisition.

### Animal preparation and 2P imaging

Larvae (5-7 dpf) were screened for *nacre-/-* and strong GCaMP expression under an epifluorescence microscope (Nikon SMZ25, Tokyo, Japan). They were mounted in a 35 mm petri dish lid in 1.6 % low melting agarose in E3. The agarose surrounding the eyes was removed to ensure unhindered eye movements [70]. During the experiment the fish were kept in E3 solution devoid of methylene blue.

### Microscope Setup

The setup was based on a previously published study [1]. In short, stimuli were presented as vertical gratings (12 roughly equally spaced, red, vertical bars per 360°) rotating horizontally around the larvae on a custom-made LED arena. Note that the 700 lp dichroic illustrated in Fig. 1b reflected only a fraction of the 850 nm IR-LED light to the sample, which still sufficed to fill out the hole in the IR-LED ring and thus provide back-illumination of the larval eyes for camera detection. Calcium signals were recorded on a hindbrain patch of ∼280 x 280 µm at 2 fps on a MOM microscope [Sutter Instruments, Novato, USA; [71]] using C7319 preamplifier (Hamamatsu Photonics K.K., Hamamatsu, Japan) and Sutter’s MScan software (Version 2.3.0.1), a 2-photon IR laser (Coherent Chameleon Vision S; 920 nm excitation wavelength; Coherent Inc., Santa Clara, USA) and a 25x Objective (Nikon CFI75, Tokyo, Japan). Stimulation and eye movement recordings were achieved via an precursory version of ZebEyeTrack [72] running in the LabVIEW environment (National instruments, Austin, USA) and a CMOS camera (DMK 23UV024, The Imaging Source GmbH, Bremen, Germany). Stimulus speed was chosen for each fish individually depending on the experiment conducted (see below) in order to preferentially generate robust slow phases covering a large dynamic range of eye positions and minimize the occurrence of quick phases (saccades).

### Stimulus protocol for the experiment on monocular versus binocular motor drive

The stimulus protocol was subdivided into three parts, each lasting for 150 seconds. In the first two parts only one eye received a moving stimulus (hereafter referred to as the stimulated eye) while the other eye received a stationary stimulus, and in the third part both eyes were stimulated. The binocular zone was blocked by black aluminium foil (BKF12, Thorlabs, Newton, USA) the whole time. Stimulus direction changed every 8-10 sec with a stable stimulus for 2-4 sec after each direction change. The average stimulus speed during motion phases across animals was 39 degrees/sec ± 11 degrees/sec (STD). Stimulus parameters were chosen for each fish individually to minimize occurrence of saccades. During monocular stimulation a stationary vertical grating was shown to the OFF eye to minimize yoking. In 137 recordings the left eye was stimulated first, in 15 the right. For illustration and analysis purposes the latter were reshaped to match the other recordings.

### Stimulus protocol for the experiment on velocity vs. position neuronal tuning

In the beginning of this stimulus protocol, an alternating OKR stimulus was presented (8 sec CW, 8 sec CCW, 12 repetitions) which was followed by a closed loop protocol in which successful completion of particular eye position/eye velocity combinations was ensured by real-time eye position monitoring. Here, eye position bins were defined, each 2° wide. In 57 recordings, bins were defined between ± 10°, in 3 recordings between ± 8°, which corresponded to the well-explored dynamic range of horizontal eye movements. For each eye position bin, the eyes were first driven via the optokinetic response into this bin and then the stimulus velocity was reduced to zero. If the larva kept its gaze centred within that bin for 4 seconds, the quality criterion was passed, and if the mean eye position moved outside the respective bin boundaries during the 4 seconds, this part was repeated until it finished successfully. Then, the eye position passed through each bin in CW and CCW directions with different stimulation speed (baseline speed, 1.2 x and 1.4 x of the baseline speed). If a saccade occurred, the current step of the protocol was repeated. The whole closed loop protocol was repeated three times. The average baseline stimulation speed was 31 degrees/sec ± 13 degrees/sec (STD). Stimulation speed was altered if fish behaviour changed during the experiment.

### Identification of neurons with oculomotor tuning (data analysis)

All data analysis was done in MATLAB (MathWorks, Natick, USA). Regions of interest (ROIs) were semi-automatically identified as previously published [Correlation Analysis, 3D mapping, [1]]. This method was altered such that we could apply several regressors at once to a recording, thus enabling us to identify neurons with different coding features at once. For this purpose, each pixel surpassing the z-score threshold for any of the regressors was coloured in the anatomical image according to its absolute maximal z-score across regressors, resulting in a heat map. This was done to identify eye movement related pixels, tighter exclusion criteria are applied later in the analysis pipeline depending on the experiment conducted. Regressors used in this study (averaged across both eyes):

- rectified low eye velocity (capped at 20 degrees/sec, separate regressors for CW and CCW directions)
- rectified high eye velocity (velocities higher than 20 degrees/sec in CW and CCW)
- angular eye position

Since the GCaMP expression was restricted to the nucleus, all drawn ROIs corresponded to somatic signals.

Each recorded optical slice was manually registered in x, y, and z planes, to a recorded z-stack of the same animal. The Mauthner cells and the medial longitudinal fasciculus (MLF) served as landmarks within the z-stack in order to combine data from multiple slices and animals into a single reference coordinate system in which the point on the midline between the Mauthner cell somata served as the origin [based on [1]]. This approach accounted for differences in the pitch, roll and yaw of individual fish. It was ignorant about inter-individual hindbrain size variations.

### Binocular coordination experiment data analysis

Data used in this experiment was recorded from 15 larvae (5-7 days post fertilization, dpf). Recordings in which the eye movements surpassed the yoking index were excluded from analysis (∼ 28 % of original recordings) beforehand (see Supplemental Fig. 1b and Supplemental Methods) which resulted in an 8-fold coverage of the imaged hindbrain region, ranging from 30 (dorsal) to −60 µm (ventral) in 5 µm intervals around the Mauthner cells (rh 4-8; xy position kept stable for different z-levels, 152 recordings total), due to previous reports of the ABN and OI location [2], [3], [12]–[14]. The oculomotor neurons of the caudal hindbrain that have been identified in this study were located mostly ventrally to the MLF stretching from the end caudal of rhombomere 6 to the ventro-caudal end of the brain. OI neurons in larval zebrafish have previously been reported ventral to the MLF and extending to the dorsal part as well [12]–[14], [46]. One study reported eye position encoding neurons in rh7/8 to be located more dorsal than other studies, but still overlapping the same volume in the brain [2]. It is therefore possible that we missed some more dorsally located OI neurons, because the dorsal parts of the hindbrain were not recorded in this study. However, an optogenetic perturbation study found the maximum effect on integrator performance in rostral areas of the OI 50 to 150 µm caudal to the Mauthner cells [17], suggesting that the relevant anatomical regions have been well sampled in this study.

To classify the response quality and type of each neuron we performed a regression analysis. For each ROI the ΔF/F (DFF) calcium time series was smoothed using a 5-time-points sliding window kernel filter, with the DFF at the time k:

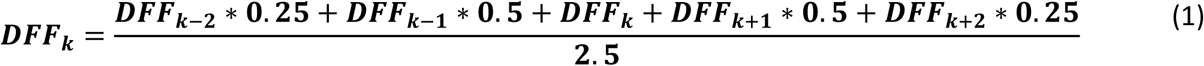

Each eye position trace was offset by its respective median to account for individual resting eye position (negative eye position and eye velocity is defined as left or leftward respectively). The DFF trace of each ROI was then correlated with several traces derived from behavioural data (eye position/velocity), which we refer to as “regressors”.

We created regressors based on conservative inclusion criteria. Each regressor was i) either coding for eye velocity or eye position, ii) had different combinations of activity during the individual stimulation phases, iii) rectified in plus or minus direction. In addition we tested two (duplicate) types of regressors sets, one in which the monocular phase activity was derived from the eye trace of the respective eye (for monocular regressors), and one in which this monocular phase activity was derived from the average of both eyes during this stimulation phase. The second set was more reliable for BA neuron identification as the motor range in the monocular phases was smaller than the one in binocular phases in most of the recordings. This resulted in a total of 52 regressors (Supplemental Fig. 1a+d).

The rectified regressors were then convolved with a “calcium impulse response function” (CIRF) [46] to account for the GCaMP dynamics in our experiments (1.1 sec measured *in vivo* by observing exponential signal decay of position encoding neurons after a saccade in the null direction). Velocity was capped at 8 degrees/sec (the regressor was set to 8 degrees/sec if the velocity exceeded 8 degrees/sec) to eliminate burst sensitivity (saccade generator). Neuronal ROIs with a correlation of at least 0.6 to any of the regressors were then kept for further analysis.

We excluded neurons from recordings in which the non-stimulated eye responded during monocular stimulus phases (Yoking index threshold, Supplemental Fig. 1b).

To exclude the possibility that some neurons were erroneously classified as monocular/binocular preferred due to eccentric firing thresholds and the fact that the dynamic eye position range differed during monocular and binocular stimulation (usually it was smaller during monocular stimulation), we calculated the firing threshold during the binocular phase and only kept neurons which reached that threshold during the monocular phases. This resulted in the exclusion of 23% (732 excluded, 2508 revised and confirmed) of neurons in this follow-up analysis (for full methods description see Supplemental Methods and Supplemental Fig. 2).

With the exception of regressors for BA neurons (r5, r6, r17, r18 for position), we did not observe any notable difference in the location or amount of identified neurons for averaged and non-averaged regressors (Supplemental Fig. 3 c, d). This is explainable by the fact that the motor range was smaller during the monocular phases and thus the resulting DFF trace is more representative of the averaged eye position trace (Supplemental Fig. 1c). As the resulting differences were small, we pooled the corresponding regressors (average and non-averaged ones) for further analysis.

### Data analysis for experiment on velocity vs. position neuronal tuning

Data used in this experiment was collected from 8 recorded fish (5-7 dpf) which resulted in a 6-fold coverage of the imaged hindbrain region (same area imaged as for binocular coordination experiment), ranging from 30 to −60 µm around the Mauthner cells in 10 µm intervals, to cover the same area as in the previous experiment (60 recordings total). ROIs were selected as previously described and considered for further analysis if their correlation to any of the rectified position or slow velocity regressors (capped at 8°/s) used in the ROI acquisition exceeded 0.4 (different threshold to previous experiment as this step was only to ensure neurons with position and velocity encoding were still included for downstream analysis). The PV_Index_ was calculated based on correlation with the respective highest scoring position and velocity regressor according to the following equation:

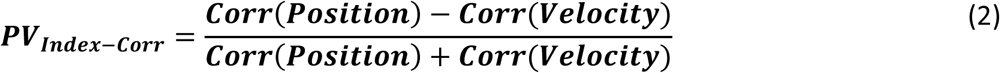

Of 889 neurons approved in the previous analysis 17 had a negative correlation for either both position or velocity regressors and were thus excluded from this PV_Index_ calculation.

For the 2 dimensional tuning curves, all frames from the recording were used (including OKR stimulation). Frames with a higher eye velocity than 10°/s and subsequent three frames were excluded to account for artefacts caused by saccades. Fluorescence was grouped in 1° eye position bins (from −15° to 15°) with the appropriate velocity (-7 degrees/sec to 7 degrees/sec) in bins of 1 degree/sec width.

### Firing threshold assessment

To extract the firing thresholds the smoothed (Eq. 1) and deconvolved (CIRF, see above) DFF was plotted against the binned eye position or velocity (2° increments for position, 1 degree/sec for velocity) tuning curve. Starting three bins from the tail (null-direction) a one sided, Bonferroni-corrected Wilcoxon rank sum test was calculated for each bin against all previous bins combined. The firing threshold was defined as the first point with significant difference to the previous (baseline) data points, where at least one of the following two bins was also significant.

To verify that inactivity of a neuron in the first experiment during a monocular stimulation phase is due to its intrinsic coding properties and not due to a lack of appropriate behaviour, the dynamic eye position range for the monocular phases was compared to the firing threshold during the binocular stimulation. If a neuron did not reach its firing threshold in any monocular phase it was excluded from further analysis (see Supplemental Fig. 2).

### Statistical information

Statistical testing was performed using MATLAB. Statistical significance level was p<0.05. For the comparison of firing thresholds in the experiment to determine the velocity and position component, a Kruskal-Wallis test was performed to check for significant differences. Other statistical tests conducted are reported in the appropriate sections.

## Declarations

### Ethics approval

All animal procedures conformed to the institutional guidelines of the University of Freiburg, University of Tübingen and local governments (Regierungspräsidium Freiburg, Regierungspräsidium Tübingen).

## Data/Material availability

Data and analysis algorithms (Matlab code) will be deposited in a public repository upon acceptance of the manuscript.

## Author contributions

C.B. and C.L. recorded the data. C.B. analysed the data. C.B., C.L. and A.B.A. wrote the manuscript. A.B.A. conceived and supervised the project and secured funding.

## Acknowledgements

We would like to thank Wolfgang Driever, Emre Aksay, Michael B. Orger, Christian Machens and Claudia Feierstein for helpful discussions on the project. Sebastian Reinig, Kun Wang, Konstantin F. Willeke and all lab members of the Arrenberg lab for discussion of the project. Bastian Hablitzel for the mapping of the abducens region in the HGj4a line. Rebecca Meier, Maximilian Wandl, Sabine Götter for excellent fish care. We would like to thank Fenja Gawlas and Philipp Rustler for help with generating transgenic zebrafish lines.

## Additional file 1

Movie1.avi: This movie shows a z-stack of a *Tg(ubi:nls-GCaMP6f)m1300* larvae at 5 dpf imaged under the above mentioned setup (except using a x20/1.0 Zeiss objective) resulting in an imaged area of 450.56 x 450.56 µm in x and y with 0.88 µm per slice in z. The movie is contrast enhanced and imaged with increased laser power (roughly 33 mW after the objective) to highlight GCaMP6f expression (same fish as in Fig. 1b).

## Supplemental information

### Supplemental Material & Methods

#### Exclusion of recordings with too much yoking

For each eye the velocity was calculated as the difference of eye position at successive time points. The eye velocity was capped at 8 degrees/sec – to prevent artefacts from saccades – and smoothed (Eq. 1). We calculated a “yoking index” (YI) according to the following equation using sums across time series data points from a given recording:

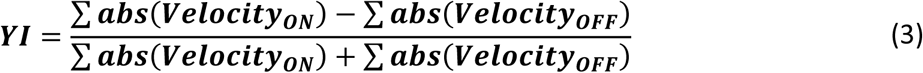

The YI was calculated for each monocular phase and only recordings where both values were bigger than 0.5 were used in the analysis. The “ON” eye was defined as the stimulated eye (Supplemental Fig. 1b).

#### Monocular coding differences (binocular coordination experiment)

For each major group of position coding neurons the correlation coefficient of the highest scoring left and right eye monocular regressor was chosen and the difference in monocular coding was calculated in the following way:

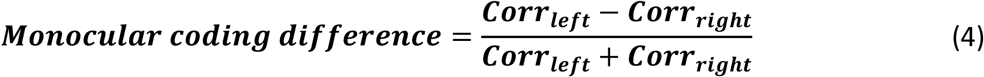

#### PV influence

For each BA and BP coding neuron the velocity influence was calculated by choosing the correlation coefficient of the appropriate velocity regressor depending on the highest scoring regressor used to identify this neuron (i.e. if the highest scoring regressor was r2 it would be compared to r10) according to:

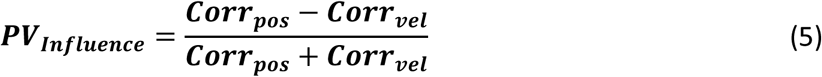

If the appropriate velocity coefficient was negative, it was set to 0.

### Chemicals and solutions

**Table 2:** Chemicals

| Chemical | Supplier: |
| --- | --- |
| NaCl | AppliChem, A3597 |
| KCl | Carl Roth, 6781.1 |
| CaCl | AppliChem, A1873 |
| MgSO4 | Merck, 1.05886.05000 |
| Methylene blue | AppliChem, A4084 |
| Agarose | Biozym, 850080 |

E3: NaCl (5 mM), KCl (0.17 mM), CaCl (0.33 mM), MgSo<sub>4</sub> (0.33 mM) with 0.01 % methylene blue.

### Supplemental Figures

**Supp. Figure 1:**
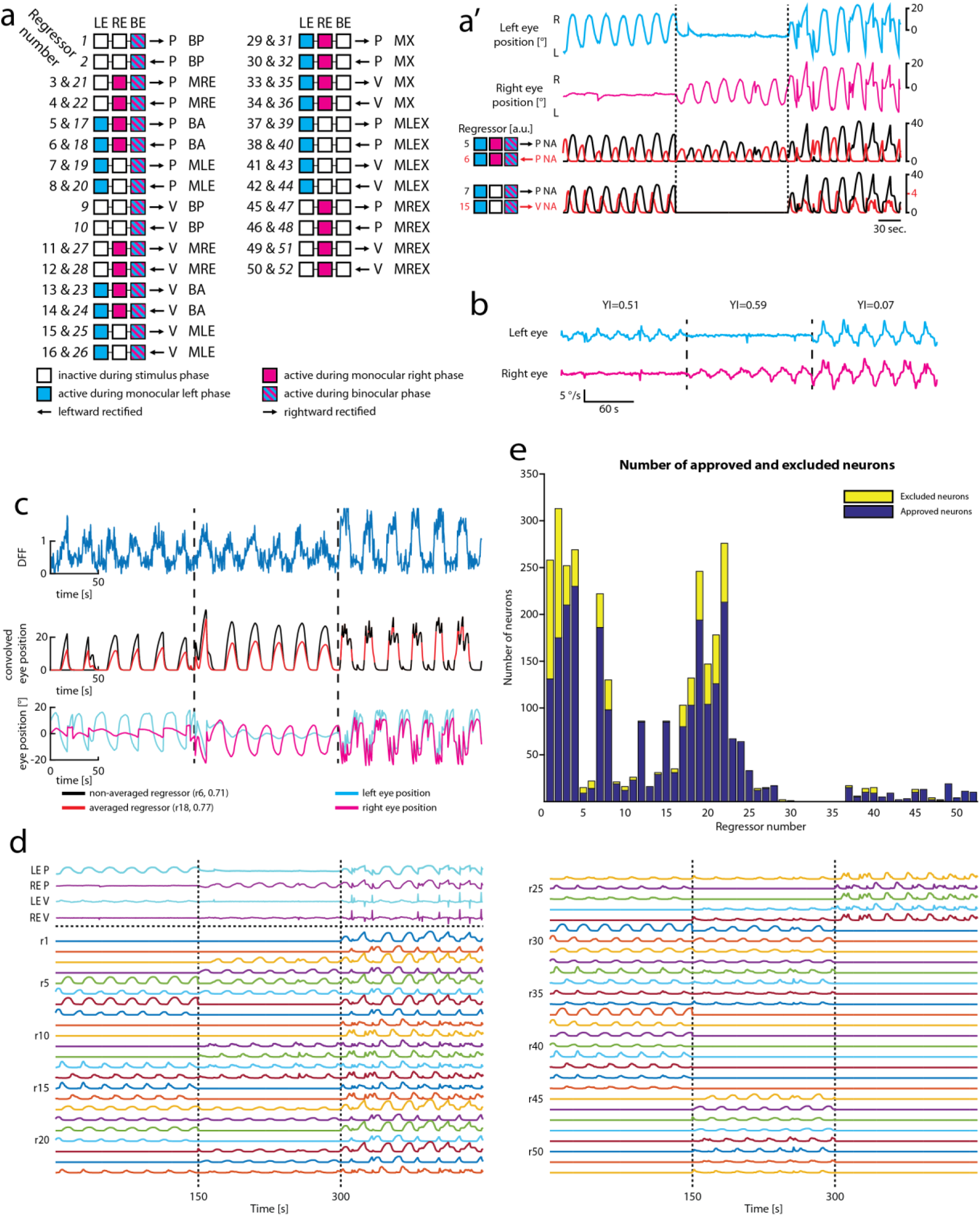
Methods for monocular/binocular analysis. **a:** Overview of regressors used to classify response types. Each set of three squares (connected by a black line) corresponds to one (or two) types of regressors, see colour legend. Regressors with italic numbers correspond to averaged regressors; BA: binocular always; BE: both eyes; BP: binocular preferred; LE: left eye; MX: monocular exclusive; MLE: monocular left eye; MLEX: monocular left eye exclusive; MRE: monocular right eye; MREX: monocular right eye exclusive; P: position; RE: right eye; V: velocity; **a’:** Example regressors and respective eye traces. NA: non-averaged (see Methods); P: position; V: velocity; eye traces same as in figure 1c-c’; **b:** Example eye traces for yoking index exclusion. YI: yoking index; **c:** Example binocular always (BA) neuron and the highest scoring regressor r6 (non-averaged) with the corresponding averaged regressor (r18) and eye traces they are based upon. **d:** All derived regressors from recording shown in figure 1c-c’. LE: left eye; P: position; RE: right eye; V: velocity; r: regressor. **e:** Overview of all approved and excluded neurons for each regressor based on the firing threshold analysis.

**Supp. Figure 2:**
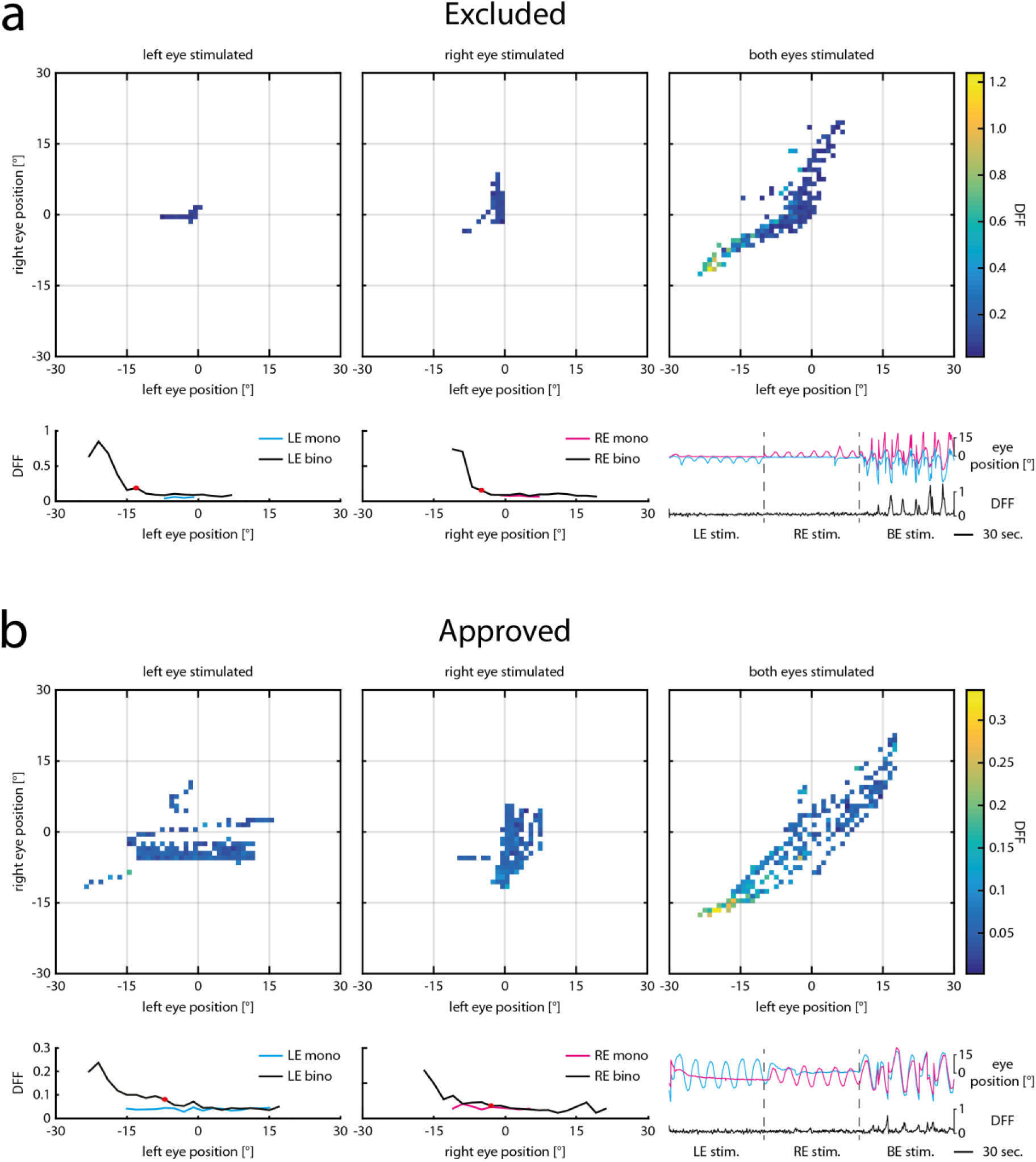
Firing threshold analysis. Tuning curves during the monocular and binocular stimulus phases for one neuron excluded from further analysis (a) and one neuron included in further analysis (b). **a:** The upper row shows the neural activity (ΔF/F) colour coded during the monocular left eye (left plot), right eye (middle plot) and binocular (right plot) stimulus phases for individual eye position bins. Monocular tuning curves (cyan left eye, magenta right eye) were plotted for the respective monocular stimulus phase and the binocular stimulus phase (black). Only bins with at least three individual data points were used. Red dot shows firing threshold. For this neuron, the eye position never explored the eye position threshold during the monocular stimulus phases, it was thus excluded from further analysis. In the lower right the corresponding eye positions and neural activity (ΔF/F) are plotted versus time. **b:** Tuning curves and eye positions for one threshold approved neuron. Note that for this neuron, the monocular tuning curves covered the eye position threshold (red dot).

**Supp. Figure 3:**
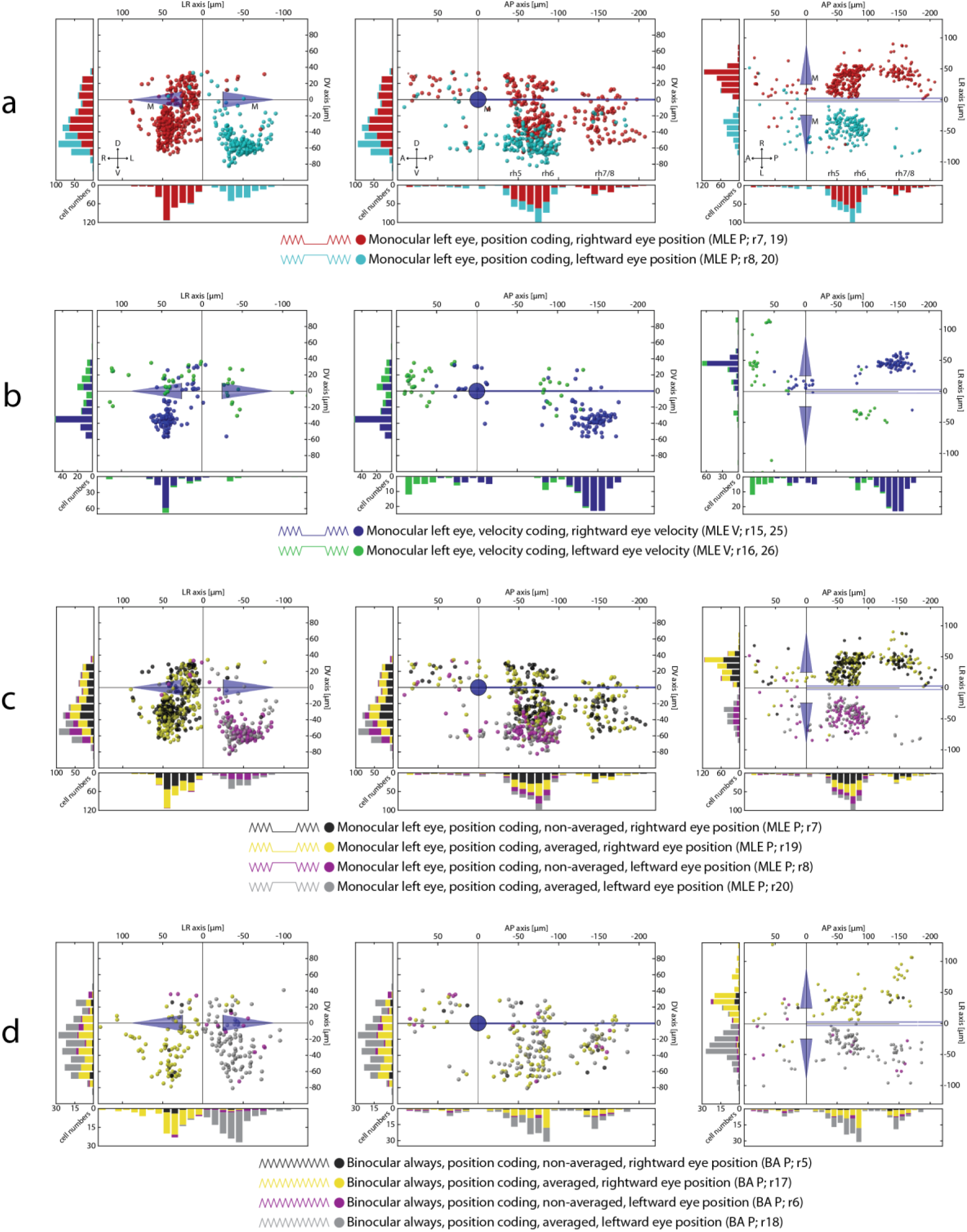
Additional monocular/binocular cell maps. **a-e:** Transversal, sagittal and dorsal views for MLE and BA neurons in the hindbrain. A: anterior; BA: binocular always; D: dorsal; L: left; M: Mauthner cells; MLE: monocular left eye; P: position/posterior; R: right; r: regressor; rh 5-8: rhombomeres 5-8; V: ventral/velocity;

**Supp. Figure 4:**
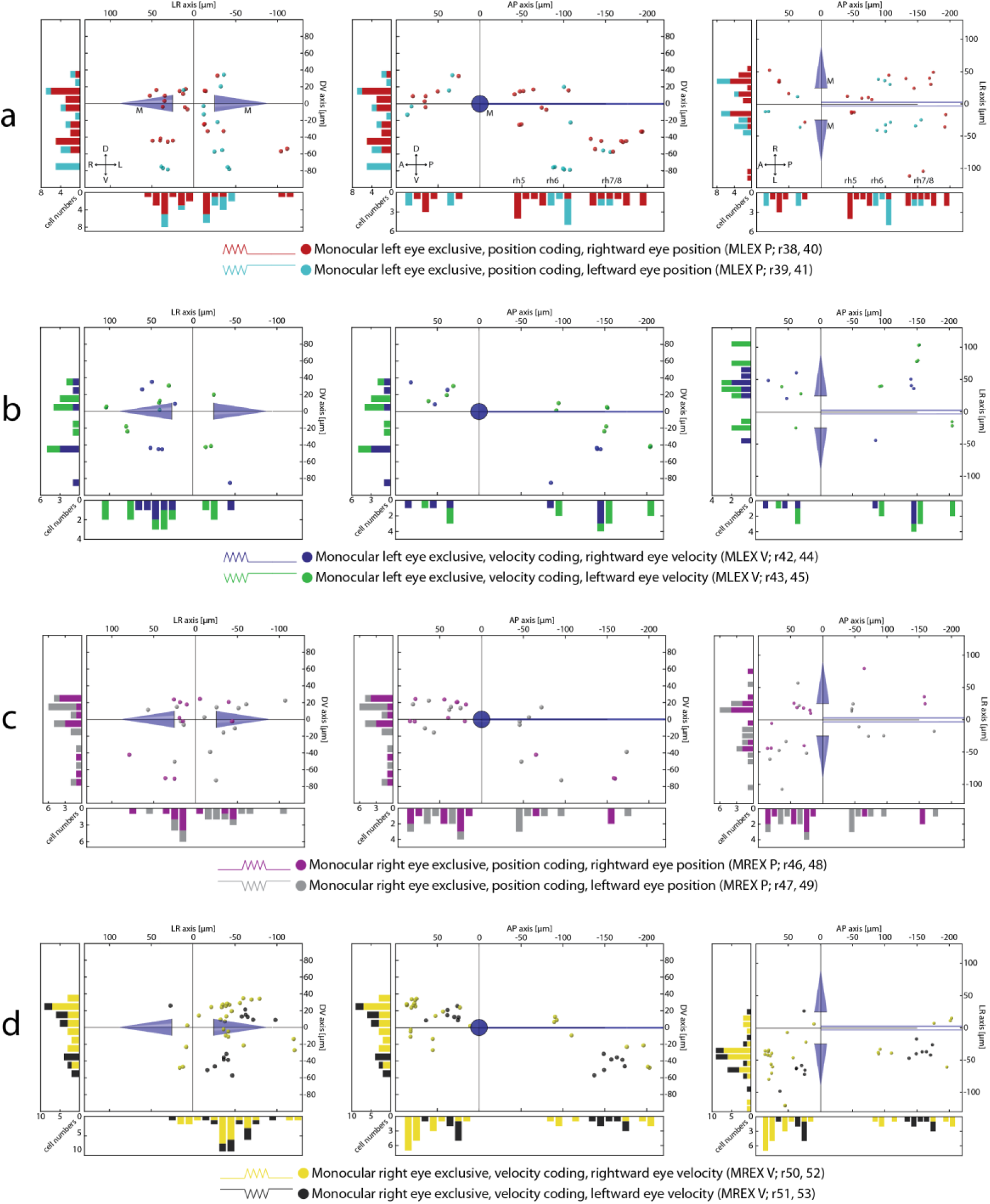
Cell maps for monocular exclusive neurons. **a-d:** Transversal, sagittal and dorsal views for MLEX and MREX neurons. A: anterior; D: dorsal; L: left; M: Mauthner cells; MLEX: monocular left eye exclusive; MREX: monocular right eye exclusive; P: position/posterior; R: right; r: regressor; rh 5-8: rhombomeres 5-8; V: ventral/velocity;

**Supp. Figure 5:**
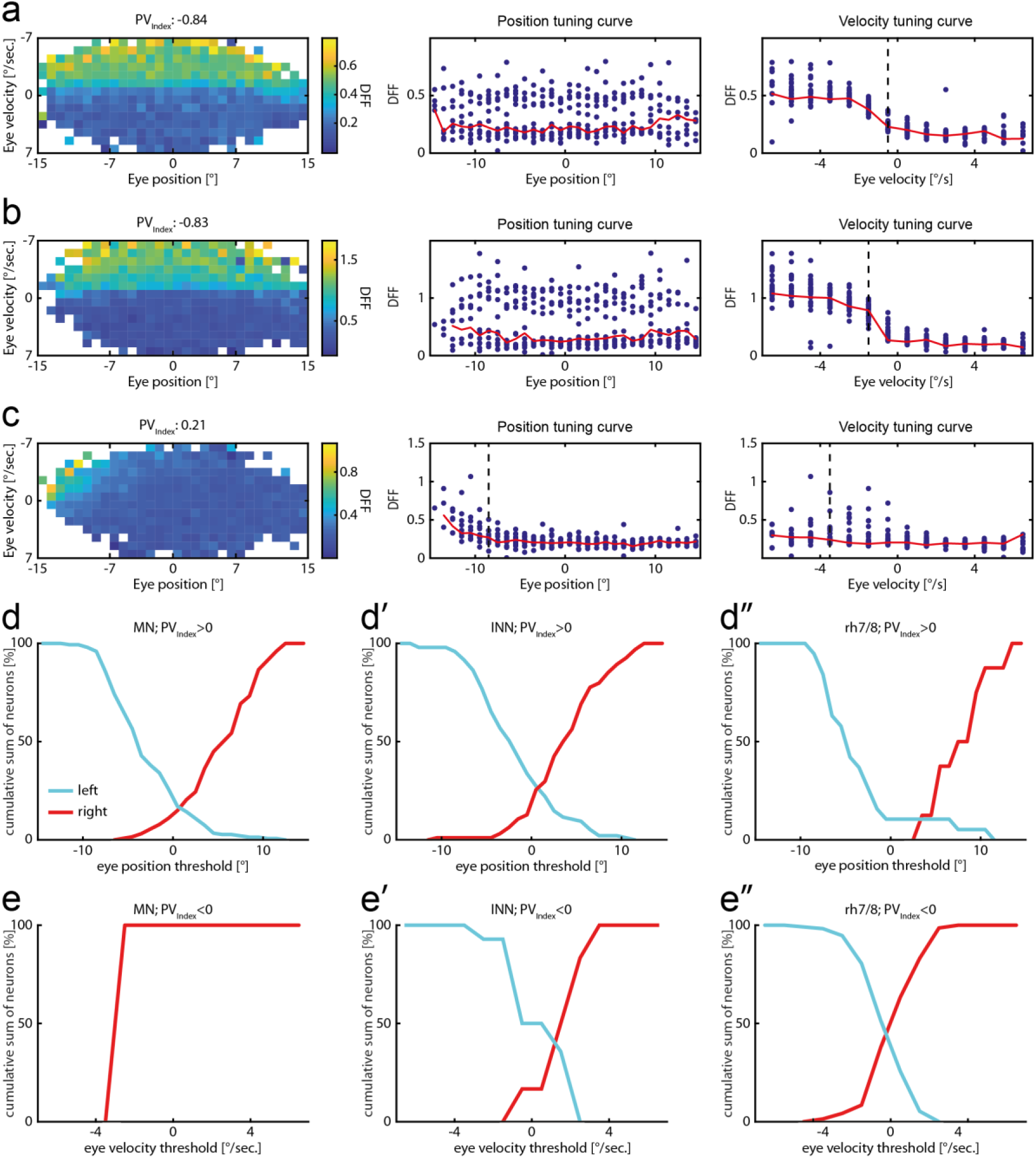
Additional tuning curves and threshold analysis. Additional tuning curves and firing thresholds for different neuron populations. **a-c:** Additional tuning curve plot same as in Fig. 4. **d-d’’:** Cumulative position threshold plots for position coding neurons (PV_Index_ > 0) pooled in ON for motoneurons (d, left: 147, right: 127), internuclear neurons (d’, left: 95, right: 94, both based on their anatomical location) and the caudal hindbrain (d’’, left: 19, right: 8). **e-e’’:** Cumulative velocity threshold plots for velocity coding neurons (PV_Index_ < 0) pooled in ON for motoneurons (e, left: 0, right: 1), internuclear neurons (e’: left: 14, right: 6) and the caudal hindbrain (e’’: left: 113, right: 71).

**Supp. Figure 6:**
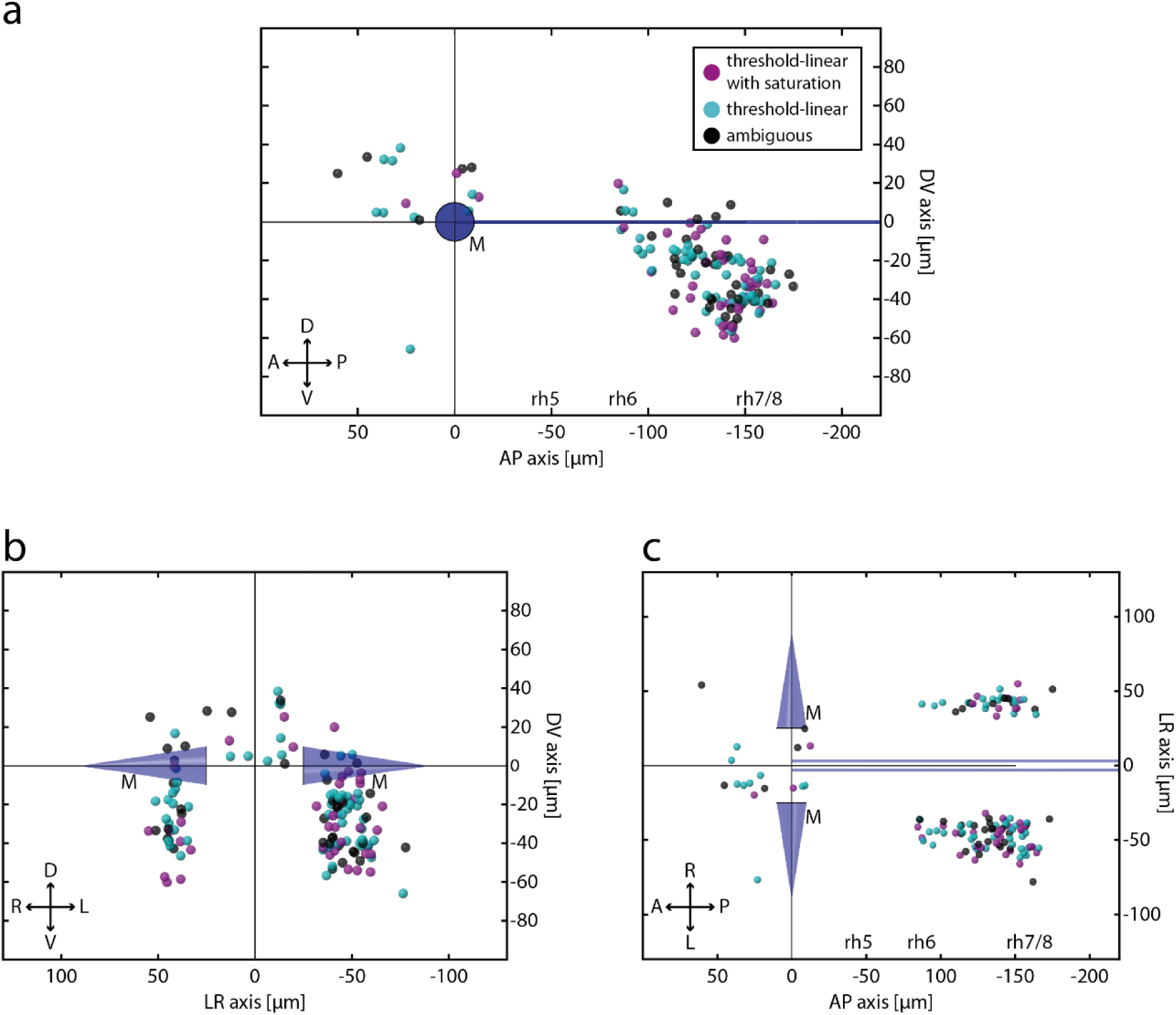
Different response profiles for velocity neurons. Velocity neurons with different response profiles show no spatial clustering. **a-c:** Sagittal, transversal and dorsal view of threshold-linear (n=60), threshold-linear with saturation (n=40) and ambiguous (n=39) neurons (PV_Index_ < −0.5) color-coded according to their response type. A: anterior; D: dorsal; L: left; M: Mauthner cells P: posterior; R: right; rh5-8: rhombomeres 5-8; V: ventral.

**Supp. Figure 7:**
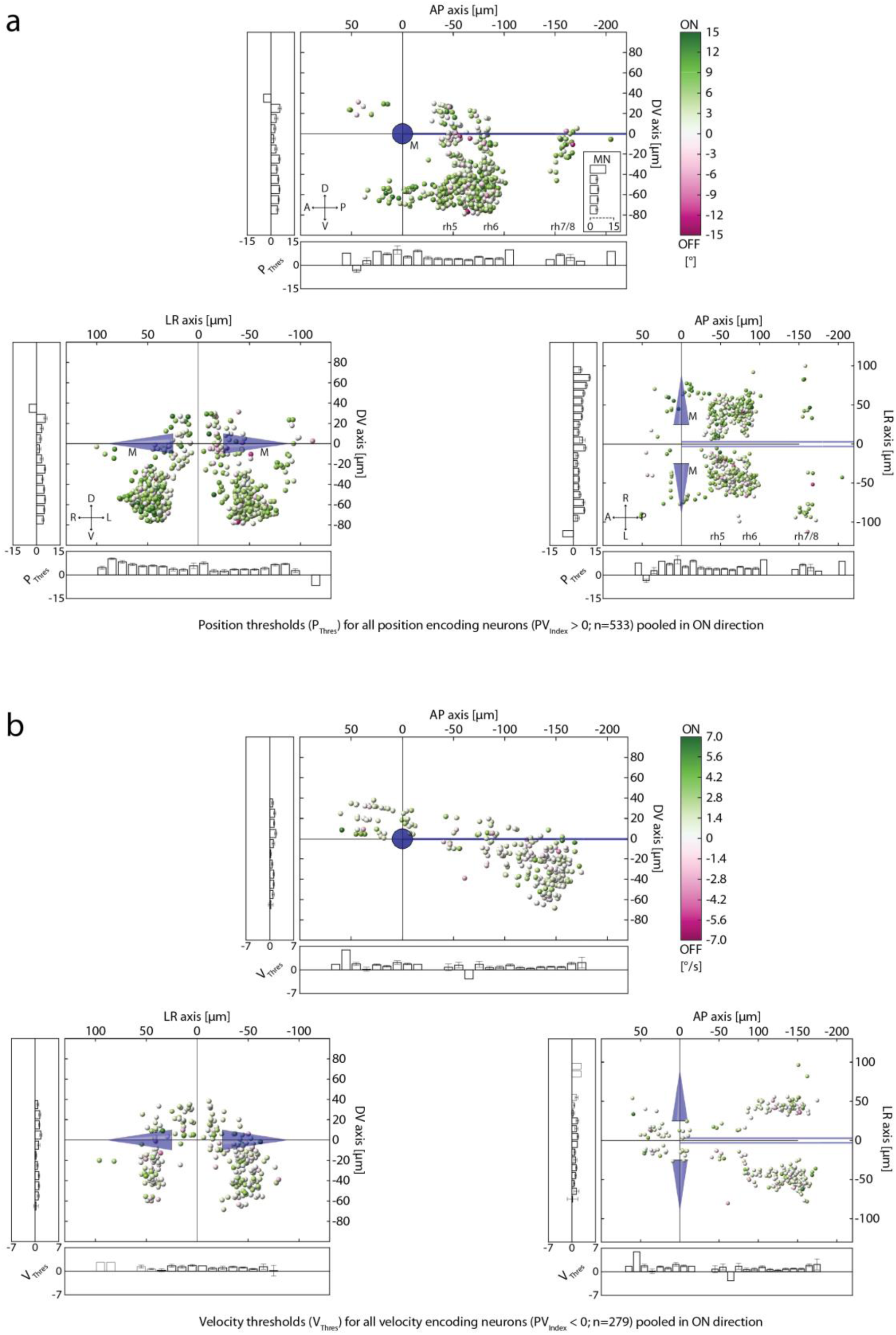
Position and velocity thresholds. Transversal, sagittal and dorsal views of position and velocity coding neurons colour-coded for their thresholds. **a:** Position thresholds (P_Thres_) colour-coded for all position coding neurons (PV_Index_ > 0) with an identified firing threshold pooled in ON direction (n=533). Inset shows thresholds for motoneurons based on their anatomical location (no statistical significance was observed: Kruskal-Wallis p=0.22; n=2, 41, 98, 89, 43) **b:** Velocity threshold (V_Thres_) colour-coded for all velocity coding neurons (PV_Index_ < 0) with an identified firing threshold pooled in ON direction (n=279).

